# P/Q-type voltage-gated calcium channels regulate calcium signaling and developmental myelination in oligodendrocyte lineage cells

**DOI:** 10.64898/2025.12.12.694003

**Authors:** Melanie Piller, Ryan A Doan, Cody L Call, Stephen M Smith, Kelly R Monk

## Abstract

Oligodendrocyte lineage cells (OLCs) are glia that arise as oligodendrocyte progenitor cells (OPCs) in the central nervous system (CNS) and may persist as progenitors or differentiate into myelin-producing oligodendrocytes (OLs). OLCs are sensitive to neuronal activity, which can influence myelin formation via activity-dependent myelination. Calcium influx in OLCs regulates many developmental processes, including stabilizing newly formed sheaths. OLCs possess P/Q-type voltage-gated calcium channels (VGCC) that contribute to calcium influx and mutations in these channels are implicated in a spectrum of neurological disorders, yet the functional significance of P/Q-type channels in OLCs is not well understood. In this study, we employ zebrafish to investigate the role of P/Q-type channels in OLCs *in vivo* during development. We use global and cell-type specific CRISPR/Cas9-mediated genome editing approaches in conjunction with live imaging and physiology to characterize the morphology and signaling properties of OLCs with mutated P/Q-type channel genes. P/Q-type channels are required for normal myelination in the developing CNS and mutants present with decreased amplitude sheath calcium transients, reduced myelin production, shorter myelin sheaths and blebbing membrane structures during development. These findings provide new insight into the role of P/Q-type calcium channels in regulating OLC development and myelination.

## Introduction

In jawed vertebrates, myelination is the process by which myelinating glia ensheathe axons and compact membranes to form an insulating layer around neurons and facilitate efficient signal propagation via saltatory conduction. In the central nervous system (CNS), myelin is formed by oligodendrocytes (OLs). OLs differentiate from OL progenitor cells (OPCs), and both these cell types are considered OL lineage cells (OLCs). OLCs must execute a complex cellular program to form myelin. Myelination requires OPC differentiation into myelinating OLs, coordination of the cytoskeleton^1^, expansion of the cell membrane^2–4^, ensheathment of target axons, stabilization and elongation of nascent sheaths^5–7^, membrane compaction, and continued remodeling during development and throughout life. This process can be modulated by many different factors. For example, differences in activity between neighboring neurons can cause OLs to preferentially myelinate more active axons through activity-dependent myelination^6,8–11^. This process is critical for learning and plasticity throughout life ^8,10–13^ and can also pathologically reinforce seizure networks in epilepsy^14^. The ways in which OLCs sense neuronal activity and how this modulates myelination are active areas of research.

Calcium signaling in OLCs is often implicated downstream of neuronal activity^5,6,15^. Calcium plays many roles in regulating cellular processes important in myelin development, including vesicle cycling, gene transcription, and cytoskeletal organization^16–20^. Calcium flows into the cytoplasm from both intracellular and extracellular sources. Extracellular calcium may enter the cell through plasma membrane ion channels, including AMPA receptors and voltage-gated calcium channels (VGCCs), which are activated by local depolarization. P/Q-type and L-type channels are VGCC subtypes present in OLCs and are activated by AMPA receptor induced local depolarization in OLCs^21,22^. Loss of L-type channels in OLCs can impair differentiation and myelination ^22,23^, but P/Q-type channels, also called Ca_v_2.1 channels, have not been thoroughly studied in OLCs. P/Q-type channels are important for neuronal signaling^24–26^ and are haploinsufficient in human CACNA1A-related disorders^27^, which include epilepsy and episodic ataxia^28–30^ and often involve white matter deficits^31–33^.

Studying how P/Q-type channel signaling impacts OLC development expands our understanding of both the basic mechanisms of developmental myelination and the role of OLCs in CACNA1A-related disorder pathology. Here, we use cell-specific and global mutagenesis strategies to study the cell-autonomous and non-autonomous effects of P/Q-type channel mutations on OLC calcium signaling and developmental myelination *in vivo* in the larval zebrafish CNS. We find that P/Q-type channel mutations cause OLs to produce less total myelin and shorter average sheaths. Using genetically encoded calcium indicators, we find that loss of P/Q-type channels reduces the amplitude of sheath calcium transient peaks and the duration of OPC calcium microdomain transients. We demonstrate that P/Q-type channel mutation results in reduced rate of sheath elongation and an increased incidence of membrane blebbing associated with sheath shortening. These results provide evidence for the role of P/Q-type channels in regulating OLC development and myelination and contribute to greater understanding of the role of calcium signaling in these cells.

## Methods

### Zebrafish husbandry/lines used

All zebrafish lines were maintained in accordance with institutional animal care standards and protocols at Oregon Health & Science University (protocol TR02_IP00001148). Experimental larvae were generated by heterozygote intercrosses of mutant background. *cacna1aa^vo105^* and *cacna1ab^vo106^* lines were maintained as heterozygotes. Larval fish were fed Gemma 150 and rotifers, and adult fish were maintained with Gemma 300 and brine shrimp. To achieve sparse labeling, we injected 50 nL of DNA at 30 ng/µL into single-cell embryos. The transgenic zebrafish lines and constructs used in this study are outlined below in Table 1.

**Table 1.** Transgenic zebrafish lines and genetic constructs used in this study.

| Stable transgenic and mutant lines | Injected DNA | Injected RNA |
| --- | --- | --- |
| *Tg(mbp:EGFP-CAAX)*<sup>34</sup> | *sox10:myrEGFP-P2A-Lifeact-TagRFP* (this study) | *CACNA1A* from pcDNA3.1<sup>35</sup> |
| *Tg(mbp-nls:EGFP)*<sup>36</sup> | *10xUAS:myrMScarlet-p2A-Cas9, U6:sgRNA*<sup>37</sup> |  |
| *Tg(olig1:myrGCaMP6s)*<sup>37</sup> |  |  |
| *Tg(sox10:KalTA4)*<sup>38</sup> |  |  |
| *Tg(10xUAS:myrGCaMP6s)*<sup>37</sup> |  |  |
| *Tg(olig1:mScarlet)*<sup>39</sup> |  |  |
| *cacnalaa*<sup>vo105</sup> (this study) |  |  |
| *cacnalab*<sup>vo106</sup> (this study) |  |  |

The pTol2-sox10:myrEGFP-P2A-Lifeact-TagRFP plasmid was generated using Gateway cloning by combining pDestTol2PA2, p5E-sox10 promoter, pME-myrEGFP, and p3E-P2A-Lifeact-TagRFP. The p3E-P2A-Lifeact-TagRFP entry vector was generated by amplifying Lifeact-TagRFP from the sox10:Lifeact-TagRFP plasmid^40^ with XhoI and SpeI overlap extension PCR primers and inserted into the p3E-P2A-MCS entry vector by restriction cloning.

### CRISPR/Cas9 mutagenesis

We used CHOPCHOP to identify potential sgRNA sequences for *cacna1aa* and *cacna1ab* CRISPR/Cas9 mutagenesis to generate the *cacna1aa^vo105^* and *cacna1ab^vo106^* stable lines. These sgRNA sequences were also used with the *10xUAS:myrmScarlet-p2A-Cas9, U6:sgRNA* constructs to induce cell-specific mutations in *cacna1aa* and *cacna1ab.* The sgRNA sequences used in this study were designed to target a locus of mutation previously found to cause loss of function resulting from single nucleotide variance in human patients^41^, with the goal of inducing loss-of-function mutations in targeted cells even if a frameshift mutation did not occur in every cell. In the human gene, this previously identified mutation is R279C, the locus of which is homologous to zebrafish *cacna1aa* amino acid 270 and *cacna1ab* amino acid 266. We used efficient sgRNAs to specifically target this locus in each gene, as shown in Table 2. The cut sites coincided with restriction sites in each gene, allowing for PCR and restriction digest to identify founders and genotype these fish. A gel demonstrating the sgRNA efficiency is shown in Supplementary Fig. 1b.

**Table 2.**
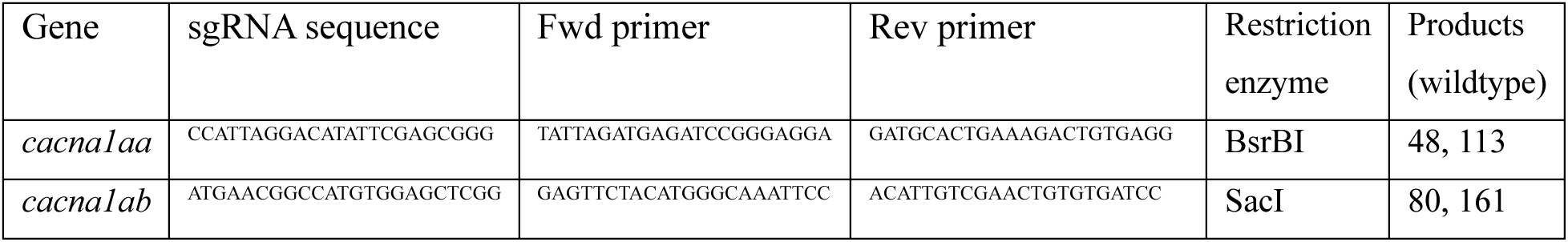
sgRNA and primer sequences for CRISPR-Cas9.

We generated stable global mutant zebrafish lines for *cacna1aa* and *cacna1ab* by injecting approximately 30 pg of the respective sgRNAs into single-cell zebrafish embryos along with 1 ng Cas9 protein in approximately 2 nL total volume. After outcrossing the initial F0 generation to wild-type AB fish, we genotyped the F1 generation to identify founders, which were then used to establish the mutant lines. The mutations selected for the *cacna1aa^vo105^* and *cacna1ab^vo106^* alleles are outlined in Table 3 and characterized in Supplementary Fig. 1.

**Table 3.** *cacna1aa* and *cacna1ab* alleles generated in this study.

| Gene | Allele | Mutation type | Predicted gene product |
| --- | --- | --- | --- |
| *cacnalaa* | *vo105* | 5 bp deletion (frameshift) | Truncated protein with 269 preserved amino acids |
| *cacnalab* | *vo106* | 7 bp deletion (frameshift) | Truncated protein with 265 preserved amino acids |

These sgRNAs were also used in conjunction with a previously published cell-type specific mutation targeting approach. We injected 30 pg of the *10xUAS:myrmScarlet-p2A-Cas9, U6:sgRNA*^37^ plasmid with sgRNA targeting either *cacna1aa* or *cacna1ab* into *Tg(sox10:KalTA)* heterozygous zygotes. This approach created sparsely labeled, mutation-targeted cells in otherwise wild-type animals. We then imaged the *myrmScarlet^+^* OLs at 3, 5, and 7 days post-fertilization (dpf) and quantified myelin sheath lengths.

### qRT-PCR

Heterozygote intercrossed embryos at 3 dpf were genotyped by tail cutting, DNA extraction, PCR, and restriction digest. RNA was extracted from genotyped pooled embryos (n = 6-12 fish per genotype) at 3 dpf with the RNeasy Mini Kit (QIAGEN) and stored at −80°C. For each genotype, cDNA was generated using SuperScript III First-Strand Synthesis (Thermo Fisher) and stored at −20°C before proceeding to qPCR with PowerUP SYBR Green (Thermo Fisher) with a QuantStudio 6 Flex Real-Time PCR System (Thermo Fisher). Primers used were as follows:

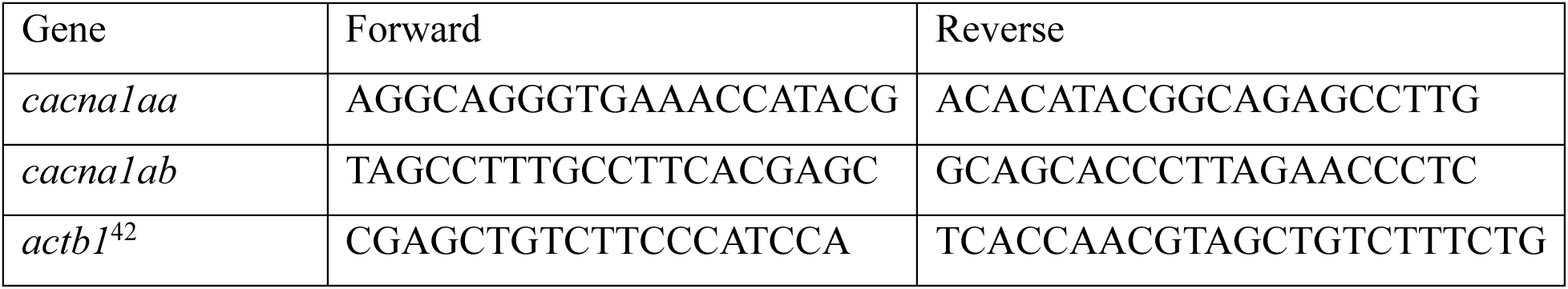

### Electrophysiology

All recordings were made in whole-cell patch-clamp in OPCs in the spinal cord of 3 dpf zebrafish embryos. The embryos were prepared by ventricle injection with TTX solution (2nL of 0.5mM TTX in 200mM KCl), then dissecting out the skin and muscle using electrosharpened tungsten needle and glass micropipette. The embryo was attached to the sylgard-lined recording chamber with electrosharpened tungsten pins and visualized using an upright microscope (Axioskop, Zeiss) while being perfused with artificial cerebrospinal fluid (in mM) 134 NaCl, 10 Glucose, 10 NaHEPES, 2.9 KCl, 2.1 CaCl_2_, 1.2 MgCl_2_ adjusted top pH 7.8 with NaOH^43^). Electrodes were positioned using a Siskiyou micromanipulator while visualized with DIC optics. OLCs were identified by expression of the mScarlet (red) fluorophore and recording configuration confirmed by the green CF488 dye. The patch electrode contained (in mM) 135 K-gluconate, 10 HEPES, 10 EGTA, 4 MgCl_2_, 4 Na_2_ATP, 0.5 NaGTP, and 0.03 CF488 dye adjusted to pH 7.2 with KOH and had a resistance of 3-5 MΏ. Voltage clamp recordings were made using an Axopatch 200B digitizer and Digidata1550B amplifier, with pClamp11 software. Voltages reported are corrected for liquid junction potentials. Images were acquired with Scicam Pro (Scientifica) camera and Ocular software.

### Imaging and Analysis

Imaging experiments were conducted using a Zeiss LSM980 confocal microscope with Airyscan2. Calcium imaging recordings were made over a 10-minute period, acquiring images at 0.5 Hz in the case of OPC calcium imaging, and 0.3-0.5 Hz in OL sheath calcium signaling depending on field size. Longer time-lapse imaging experiments (see Figures 6 and 7, Supplementary videos 3 and 4) were conducted at 1 image per 15 minutes. Images were processed using Zeiss Airyscan processing, then exported to ImageJ for quantification and analysis. Calcium imaging analysis was performed by using a semi-supervised ImageJ ROI selection process to identify calcium microdomains using the GECIquant plugin^44^. We then measured the fluorescence intensity in the ROI over the course of the time-lapse, and used MATLAB to identify peak events, defined as increases in fluorescence intensity over baseline greater than three times the standard deviation, and quantify characteristics of these peaks including amplitude and duration. For more details, see these methods as previously described^37^. Myelin sheath quantification was made by tracing and measuring individual sheath lengths in ImageJ, as previously described^37^. Lifeact signal analysis was performed with cells that had at least two hours of imaging data after differentiation (i.e. after all OPC processes had disappeared), by tracing the bottom of a sheath using the membrane fluorophore and the Lifeact signal using simple neurite tracer^45^. Then, using a custom MATLAB script, we measured the distance between the Lifeact signal and the bottom of the sheath across time-lapse imaging (for 2 sheaths and 2-6 hours per cell) and quantified the sum absolute value of changes in this distance during the measured period, averaged along the length of the sheath to get an estimate of variability in Lifeact signal per hour. Manipulations of example images were performed in ImageJ and limited to adjusting brightness, rotating, and cropping images.

### Transmission electron microscopy

Zebrafish larvae were anesthetized with tricaine and sectioned between segments 5 and 7 to standardize sampling along the anterior-posterior axis. Tissues were fixed in a modified Karnovsky’s solution (2% glutaraldehyde, 4% paraformaldehyde in 0.1 M sodium cacodylate buffer, pH 7.4; all chemicals from Electron Microscopy Sciences). Samples were microwaved (PELCO BioWave, Ted Pella) using a sequence of 100 W for 1 min, off for 1 min, 100 W for 1 min, off for 1 min, followed by 450 W for 20 seconds, and off for 20 seconds, repeated five times. Samples were then fixed overnight at 4°C. The next day, samples were rinsed three times in 0.1 M sodium cacodylate buffer at room temperature, with each rinse lasting 10 minutes. A secondary fixation was performed using a 2% osmium tetroxide solution, prepared by mixing 2 mL of 0.2 M sodium cacodylate with 0.2 M imidazole (pH 7.5) and 2 mL of 4% osmium tetroxide (Electron Microscopy Sciences). Samples were treated with this solution and microwaved using the same sequence as above (100 W for 1 min, off for 1 min, 100 W for 1 min, off for 1 min, 450 W for 20 seconds, off for 20 seconds, repeated five times). They were then incubated at room temperature for 2 hours. The osmium tetroxide was removed, and samples were washed three times in deionized water, 10 minutes per wash. Next, UranyLess (Electron Microscopy Sciences) was added, and samples were microwaved at 450 W for 1 min, off for 1 min, and 450 W for 1 min. The UranyLess was removed, and samples were washed three times in deionized water, 10 minutes each. Samples were dehydrated using a graded ethanol series (25%, 50%, 70%, 80%, 95%, and 100% ethanol in water), with a pause at the 70% ethanol step, leaving the samples in it overnight at 4°C before continuing with 80% ethanol the next day. For each ethanol step, samples were microwaved at 250 W for 45 seconds, followed by a 10-minute incubation at room temperature. For the 100% ethanol step, samples were microwaved at 250 W for 1 minute, off for 1 minute, and 250 W for 1 minute, then incubated at room temperature for 10 minutes; this was repeated three times. Subsequently, samples were dehydrated in 100% EM-grade acetone, microwaved at 250 W for 1 minute, off for 1 minute, and 250 W for 1 minute, followed by a 10-minute incubation at room temperature. This acetone step was repeated three times. For embedding, samples were infiltrated with a 1:1 mixture of Araldite 812 and 100% acetone at room temperature overnight. The next day, samples were transferred to fresh 100% Araldite 812. Using a stereomicroscope (Zeiss Stemi 508), samples were carefully positioned in molds to ensure proper orientation for sectioning. The resin was allowed to set at room temperature for 4–6 hours before polymerization in a 65°C oven for at least 48 hours. For all samples, semithin sections (250 nm) were prepared and stained with 1% toluidine blue (Fisher Scientific). These sections were examined under a light microscope (Leica DM 300) to confirm sample quality before transmission electron microscopy (TEM) preparation. Ultrathin sections (70 nm) were then cut and mounted on 2×1 mm Formvar-coated copper slot grids (Electron Microscopy Sciences). The sections were counterstained with UranyLess (Electron Microscopy Sciences) and 3% lead citrate (Electron Microscopy Sciences). Images were captured using an FEI Tecnai T12 TEM microscope equipped with an Advanced Microscopy Techniques CCD camera.

We used these TEM images to compare the number of myelinated axons in the ventral spinal cord across sample groups. To quantify this, we identified the ventral medial axon tract in each image, here defined as the axons medial to the Mauthner axon on one side of the spinal cord. In this region, we counted the axons with myelin (i.e. with electron-dense membrane wraps) and compared this metric across groups.

### Quantification and Statistics

All images were analyzed under blinded conditions. In heterozygote intercross experiments, all analyses were performed prior to genotyping. In cell-specific CRISPR/Cas9 experiments, a lab member recorded sample identities, the de-identified animals were imaged and analyzed, and the samples were subsequently re-identified. Whenever possible, we used sibling controls for all comparisons with experimental groups. We used Prism to generate graphs and perform statistical tests. Briefly, multi-group analyses were performed with one-way ANOVA with Tukey’s multiple comparison post-test of all groups at a significance threshold of α = 0.05. Significant comparisons are shown in the figure graphs, and all ANOVA and post-test results are listed in the figure legend. The Brown-Forsythe test and Bartlett’s test were used to assess differences in variance among groups and validate the use of one-way ANOVA in these samples. Unless otherwise indicated, these tests do not show significant differences in the variance across groups. Unless otherwise specified, the bar plot and error bars represent the mean and standard deviation in all graphs. Unfilled data points correspond to representative images used in figures. For all data, the sample size and statistical tests are reported in each figure legend. Statistics were computed using per-animal or per-cell averages, depending on the experiment and denoted in the figure legend. P-values for all comparisons significant at the α = 0.05 level are listed in figure legends.

## Results

### OLCs express genes encoding P/Q-type voltage-gated calcium channels, and global mutation reduces expression of these genes

*Cacna1a*, the gene which encodes the α-subunit of P/Q-type calcium channels in mammals, is duplicated into two paralogs in zebrafish: *cacn1aa* and *cacna1ab*. Prior transcriptomic analyses show that zebrafish spinal cord OLCs express both *cacna1aa* and *cacna1ab* at 3 days post-fertilization (dpf) (Supplementary Fig. 1a)^46,47^. Using CRISPR/Cas9-mediated mutagenesis (Table 2; Supplementary Fig. 1b), we generated global mutant zebrafish lines possessing alleles with a premature stop codon for each gene: *cacna1aa^vo105^* and *cacna1ab^vo106^*. The resultant alleles have substantially truncated predicted gene products with 269 and 265 amino acids preserved in *cacna1aa and cacna1ab* respectively (compared to 2265 and 2285 amino acids in the wild-type proteins; Supplementary Fig. 1c). We used RT-qPCR to measure the expression of each gene in the mutant zebrafish embryos at 3 dpf and found an approximately 30% reduction of *cacna1aa* expression in the *cacna1aa^vo105/vo105^* animals, and a near complete ablation of expression of *cacna1ab* expression in the *cacna1ab^vo106/vo106^* animals (Supplementary Fig. 1d, e, ANOVA p = 0.0006 and p < 0.0001, respectively). The gross morphology of the homozygous and heterozygous *cacna1aa^vo105^* and *cacna1ab^vo106^* zebrafish larvae is largely similar to the wild-type larvae at 5 dpf, though some larvae did not have visible swim bladders by 5 dpf (Supplementary Fig. 1f, g, Chi-square p = 0.1136). Using these newly generated global mutant zebrafish lines, we next explored how P/Q-type channel mutation affects developmental myelination.

### OL P/Q-type channels regulate the length of developing myelin sheaths in a cell-autonomous manner

Myelin properties including sheath length and thickness can regulate action potential propagation efficiency, and abnormal sheath length can impede normal signaling^6,8,9^. To test if P/Q-type channels mediate development of myelin sheaths *in vivo*, we measured the length of myelin sheaths in sparsely labeled OLs in global mutant or wild-type animals at 3, 5, and 7 dpf (Fig. 1a, b). To do this, we intercrossed heterozygous *cacna1aa^vo105/+^* or *cacna1ab^vo106/+^* zebrafish, then injected the offspring at the one-cell stage with a *sox10:myrEGFP* genetic construct to sparsely label OLCs and quantified average sheath length and total length of sheaths produced for each cell across development (Fig. 1c-h). OLs at 5 and 7 dpf were initially imaged at 3 dpf to ensure they had differentiated into OLs by 3 dpf. We found the *cacna1aa^vo105^* and *cacna1ab^vo106^* heterozygous and homozygous mutant (i.e., *cacna1aa^vo105/+^*, *cacna1aa^vo105/vo105^*, *cacna1ab^vo106/+^*, and *cacna1ab^vo106/vo106^*) OLs produced shorter average myelin sheaths than the corresponding wild-type cells at 3, 5, and 7 dpf, with the exceptions of *cacna1aa^vo105/vo105^* cells at 5 dpf and *cacna1ab^vo106/+^*cells at 7 dpf, which showed a similar trend (Fig. 1c, e, g, 3 dpf ANOVA p < 0.0001 for both genotype groups, 5 dpf ANOVA p = 0.0121 for *cacna1aa*, p < 0.0001 for *cacna1ab*, 7 dpf ANOVA p < 0.0001 for *cacna1aa*, p = 0.0065 for *cacna1ab*). We also compared the total length of myelin produced by each cell and found the *cacna1aa^vo105/+^* and *cacna1aa^vo105/vo105^* OLs had a reduced total myelin output compared to sibling wild-type OLs at 5 and 7 dpf, while the *cacna1ab^vo106/vo106^* OLs produced less total myelin compared with the wild-type OLs at 5 dpf and similar trends at 7 dpf (Fig 1f, h, 5 dpf ANOVA p = 0.0083 for *cacna1aa*, p = 0.0154 for *cacna1ab*, 7 dpf ANOVA p < 0.0001 for *cacna1aa* and p = 0.1098 for *cacna1ab*), but not at 3 dpf (Fig. 1d, ANOVA p = 0.4064 for *cacna1aa*, p = 0.2478 for *cacna1ab*). These results suggest P/Q-type channels are necessary for regulating myelin sheath length and likely play a role in modulating total myelinating capacity during development.

**Figure 1.**
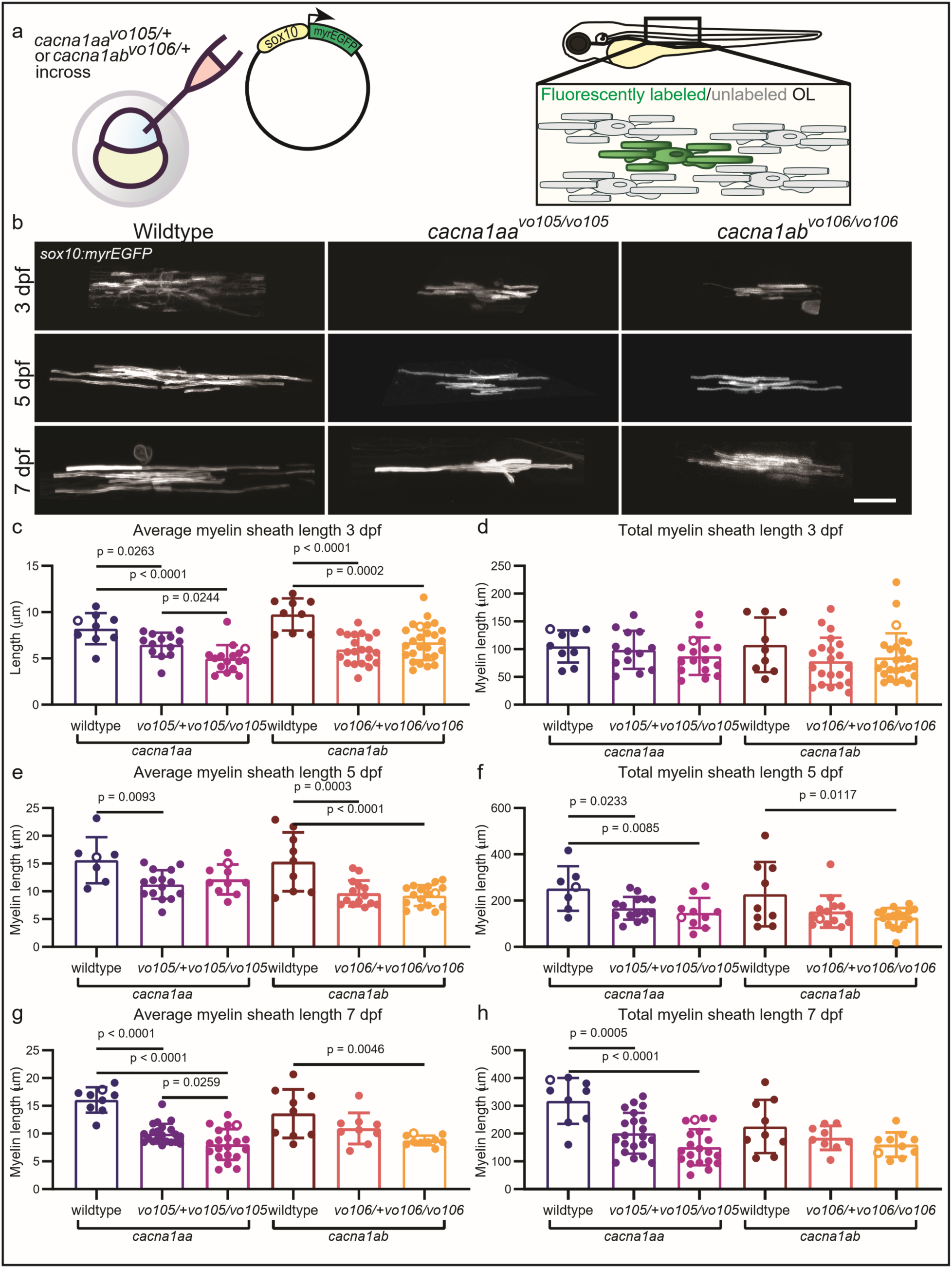
Global P/Q-type channel mutation results in reduced total OL myelin output, reduced sheath length. a. Schematic of experimental setup: *sox10:myrEGFP* DNA construct injected into single-cell embryos from heterozygous intercross of *cacna1aa^vo105/+^* or *cacna1ab^vo106/+^* to sparsely label OLs. b. Example images of sparsely labeled OLs imaged *in vivo* in zebrafish spinal cord at 3, 5, and 7 dpf. c. Average myelin sheath length across genotypes at 3 dpf (*cacna1aa* ANOVA p < 0.0001, Tukey’s multiple comparison wildtype vs *cacna1aa^vo105/+^*p = 0.0263, wildtype vs *cacna1aa^vo105/vo105^* p < 0.0001, *cacna1aa^vo105/+^* vs *cacna1aa^vo105/vo105^* p = 0.0244. *cacna1ab* ANOVA p < 0.0001, wildtype vs *cacna1ab^vo106/+^* p < 0.0001, wildtype vs *cacna1ab^vo106/vo106^* p = 0.0002). d. Total length of myelin produced by each OL at 3 dpf (*cacna1aa* ANOVA p = 0.4064, *cacna1ab* ANOVA p = 0.2478). e. Average myelin sheath length across genotypes at 5 dpf (*cacna1aa* ANOVA p = 0.0121, Tukey’s multiple comparisons wildtype vs *cacna1aa^vo105/+^*p = 0.0093, wildtype vs *cacna1aa^vo105/vo105^* p = 0.0664, *cacna1ab* ANOVA p < 0.0001, wildtype vs *cacna1ab^vo106/+^* p = 0.0003, wildtype vs *cacna1ab^vo106/vo106^* p < 0.0001). f. Total myelin sheath length per cell across genotypes at 5 dpf (*cacna1aa* ANOVA p = 0.0083, Tukey’s multiple comparisons test wildtype vs *cacna1aa^vo105/+^* p = 0.0233, wildtype vs *cacna1aa^vo105/vo105^* p = 0.0085, *cacna1ab* ANOVA p = 0.0154 wildtype vs *cacna1ab^vo106/+^* p = 0.0882, wildtype vs *cacna1ab^vo106/vo106^* p = 0.0117). g. Average myelin sheath length across genotypes at 7 dpf (*cacna1aa* ANOVA p < 0.0001, Tukey’s multiple comparisons wildtype vs *cacna1aa^vo105/+^*p < 0.0001, wildtype vs *cacna1aa^vo105/vo105^* p < 0.0001, *cacna1aa^vo105/+^* vs *cacna1aa^vo105/vo105^* p = 0.0259, *cacna1ab* ANOVA p = 0.0065, wildtype vs *cacna1ab^vo106/+^* p = 0.1614, wildtype vs *cacna1ab^vo106/vo106^* p = 0.0046). h. Total myelin sheath length per cell across genotypes at 7 dpf (*cacna1aa* ANOVA p < 0.0001, Tukey’s multiple comparison wildtype vs *cacna1aa^vo105/+^* p = 0.0005, wildtype vs *cacna1aa^vo105/vo105^* p < 0.0001, *cacna1aa^vo105/+^*vs *cacna1aa^vo105/vo105^* p = 0.0744, *cacna1ab* ANOVA p = 0.1098). Points on graph represent per-cell averages, with unfilled points corresponding to representative images in b. Sample size for each group is as follows. 3 dpf: *cacna1aa* wildtype N = 5 fish, 9 cells, *cacna1aa^vo105/+^*N = 8 fish, 13 cells, *cacna1aa^vo105/vo105^* N = 7 fish, 16 cells, *cacna1ab* wildtype N = 6 fish, 9 cells, *cacna1ab^vo106/+^*N = 11 fish, 21 cells, *cacna1ab^vo106/106^* N = 13 fish, 26 cells. 5 dpf: *cacna1aa* wildtype N = 4 fish, 7 cells, *cacna1aa^vo105/+^*N = 7 fish 15 cells, *cacna1aa^vo105/vo105^* N = 4 fish, 10 cells, *cacna1ab* wildtype N = 6 fish, 9 cells, *cacna1ab^vo106/+^*N = 7 fish, 14 cells, *cacna1ab^vo106/106^* N = 6 fish, 17 cells. 7 dpf: *cacna1aa* wildtype N = 5 fish, 9 cells, *cacna1aa^vo105/+^*N = 9 fish, 21 cells, *cacna1aa^vo105/vo105^* N = 9 fish, 20 cells, *cacna1ab* wildtype N = 6 fish, 9 cells, *cacna1ab^vo106/+^* N = 6 fish, 9 cells, *cacna1ab^vo106/106^* N = 4 fish, 10 cells.

To further examine myelin in P/Q-type channel mutants, we also performed ultrastructure analysis with transmission electron microscopy (TEM) on 5 dpf zebrafish spinal cords (Supplementary Fig. 2a). We found there were fewer myelinated axons in the ventral medial spinal cord in *cacna1ab^vo106/vo106^* mutant zebrafish compared to wildtype, and a similar trend in *cacna1aa^vo105/vo105^* zebrafish (Supplementary Fig. 2b, ANOVA p = 0.0896 for *cacna1aa,* p = 0.0264 for *cacna1ab*). Together with the sparse labeling experiment, these results demonstrate that the global P/Q-type channel mutants have impaired myelination during development. However, in this global mutant approach, changes to neuronal signaling due to mutant P/Q-type channels may also contribute to this phenotype.

To determine if there is a cell-autonomous component to P/Q-type channel regulation of OL myelin development, we next used a previously published cell-specific CRISPR/Cas9 system^37^ to sparsely label OLs targeted for mutation, then performed live imaging in the zebrafish spinal cord at 3, 5, and 7 dpf (Fig. 2a). The injected DNA constructs encoded Cas9, membrane-tethered mScarlet, and either sgRNA targeting the gene of interest or a random control sequence (Fig. 2a). This approach allowed us to measure myelin sheath length in mutated cells in an otherwise wild-type animal. We measured lengths of individual sheaths from the sparsely labeled cell-specific mutant and control OLs (Fig. 2b) and found the *cacna1aa* and *cacna1ab* targeted OLs produced shorter average myelin sheaths than their control counterparts at 3, 5, and 7 dpf. (Fig. 2c, ANOVA < 0.0001 for each timepoint). The targeted cells also produced less total myelin, i.e. sum of all sheath lengths per cell at 3 and 5 dpf (Fig. 2d, ANOVA p < 0.0001 for 3 dpf and p = 0.0020 for 5 dpf). In both the global and cell-specific mutant approaches, the wild-type or control OLs had a similar number of sheaths per OL as the mutant OLs across all timepoints (Supplementary Fig. 3a-d). The shorter average myelin sheaths and reduced myelin production by OLs with global or cell-specific disrupted P/Q-type channels suggests a cell-intrinsic signaling function of these channels in promoting normal myelin development.

**Figure 2.**
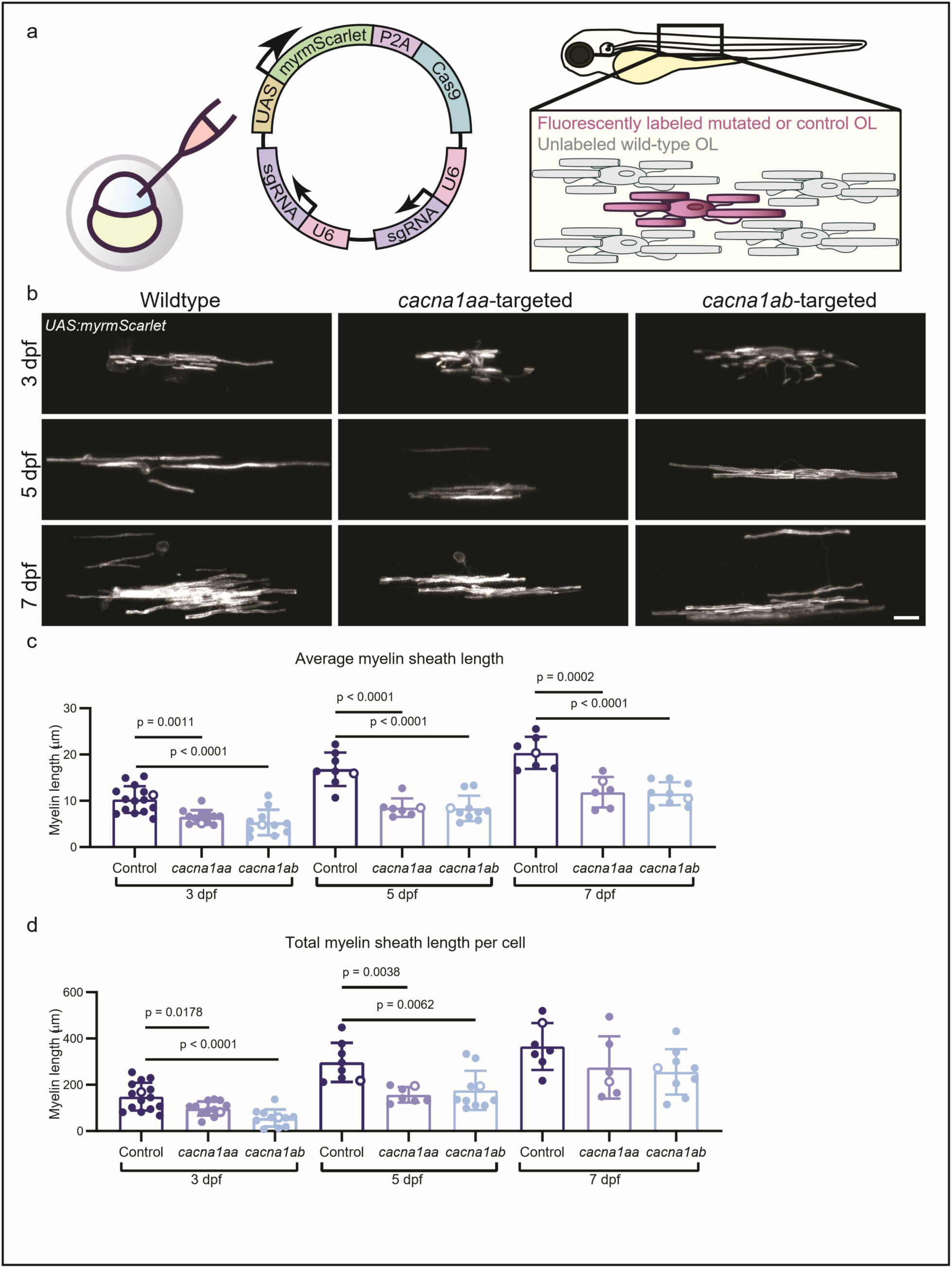
Cell-specific mutation of P/Q-type channel genes in OLs results in shorter myelin sheaths and reduced total myelin output. a. Schematic of experimental setup. b. Example images of cell-specific mutation-targeted OLs imaged *in vivo* in zebrafish spinal cord at 3, 5, and 7 dpf. c. Average myelin sheath length across conditions and timepoints (3 dpf ANOVA p < 0.0001, Tukey’s multiple comparisons test control vs *cacna1aa* p = 0.0011, 3 dpf control vs *cacna1ab* p < 0.0001, 5 dpf ANOVA p < 0.0001, control vs *cacna1aa* p < 0.0001, control vs *cacna1ab* p < 0.0001, 7 dpf ANOVA p < 0.0001, control vs *cacna1aa* p = 0.0002, control vs *cacna1ab* p < 0.0001). d. Total myelin sheath length per cell across conditions and timepoints (3 dpf ANOVA p < 0.0001, Tukey’s multiple comparisons test control vs *cacna1aa* p = 0.0178, control vs *cacna1ab* p < 0.0001, 5 dpf ANOVA p = 0.0020 control vs *cacna1aa* p = 0.0038, control vs *cacna1ab* p = 0.0062, 7 dpf ANOVA p = 0.1493). Points on graph represent per-cell averages, with unfilled points corresponding to representative images in b. Sample sizes are as follows. 3 dpf Control N = 10 fish, 15 cells, *cacna1aa* N = 7 fish, 12 cells, *cacna1ab* N = 7 fish, 11 cells, 5 dpf Control N = 6 fish, 8 cells, *cacna1aa* N = 5 fish, 7 cells, *cacna1ab* N = 5 fish, 10 cells. 7 dpf Control N = 5 fish, 7 cells, *cacna1aa* N = 4 fish, 6 cells, *cacna1ab* N = 4 fish, 9 cells.)

### P/Q-type channel mutation reduces calcium transient amplitude in developing myelin sheaths and calcium transient duration in OPC microdomains

Previous studies have implicated calcium transient activity in stabilizing and promoting elongation of developing myelin sheaths^5,6,48^. To test if the mechanisms underlying the regulation of developmental myelin sheath length involve P/Q-type channels, we measured calcium signals in the developing sheaths using a sparse transgenic membrane-tethered calcium indicator, *Tg(sox10:KalTA;UAS:myrGCaMP6s)*. For this genetically encoded calcium indicator fluorescence is proportional to intracellular calcium concentration, and it has previously been used to study myelin sheath calcium transients during development^5,6^. We recorded endogenous calcium activity over a ten-minute period, imaging at 0.5 Hz (Fig. 3a, b, Supplementary Video 1). We next measured individual sheath activity (Fig. 3c) and used a previously published method^37^ to identify and characterize calcium transient peaks, defined as deviations from baseline greater than three standard deviations. We then compared calcium peak amplitude and duration (Fig. 3d-g). Based on previous research showing activation of VGCCs in OLCs downstream of AMPAR activation^21^, we hypothesized that the amplitude and/or duration of the calcium peaks would be reduced in the *cacna1aa^vo105/vo105^* and *cacna1ab^vo106/vo106^* OL sheaths compared to the wild-type OL sheaths. In accordance with our hypothesis, we found a reduced amplitude of calcium peaks in the *cacna1aa^vo105/+^, cacna1aa^vo105/vo105^*, and *cacna1ab^vo106vo106^* fish compared to wild-type siblings at both 3 and 5 dpf (Fig. 3d, *cacna1aa* ANOVA p = 0.0072, *cacna1ab* ANOVA p = 0.0167, Fig. 3f, *cacna1aa* ANOVA p = 0.0096, *cacna1ab* ANOVA p = 0.0021). However, the average duration of sheath calcium peaks was similar across genotypes at both timepoints (Fig. 3e, *cacna1aa* ANOVA p = 0.8627, *cacna1ab* ANOVA p = 0.4760, Fig. 3g *cacna1aa* ANOVA p = 0.8821, *cacna1ab* ANOVA p = 0.7540). The frequency of peak events in sheaths with calcium activity was also similar across genotypes at both 3 and 5 dpf (Supplementary Fig. 4a, *cacna1aa* ANOVA p = 0.7024, *cacna1ab* ANOVA p = 0.3287, Supplementary Fig. 4b, *cacna1aa* ANOVA p = 0.8428, *cacna1ab* ANOVA p = 0.9819). These results suggest that calcium influx via P/Q-type channels modulates the amplitude of calcium activity in developing myelin sheaths.

**Figure 3.**
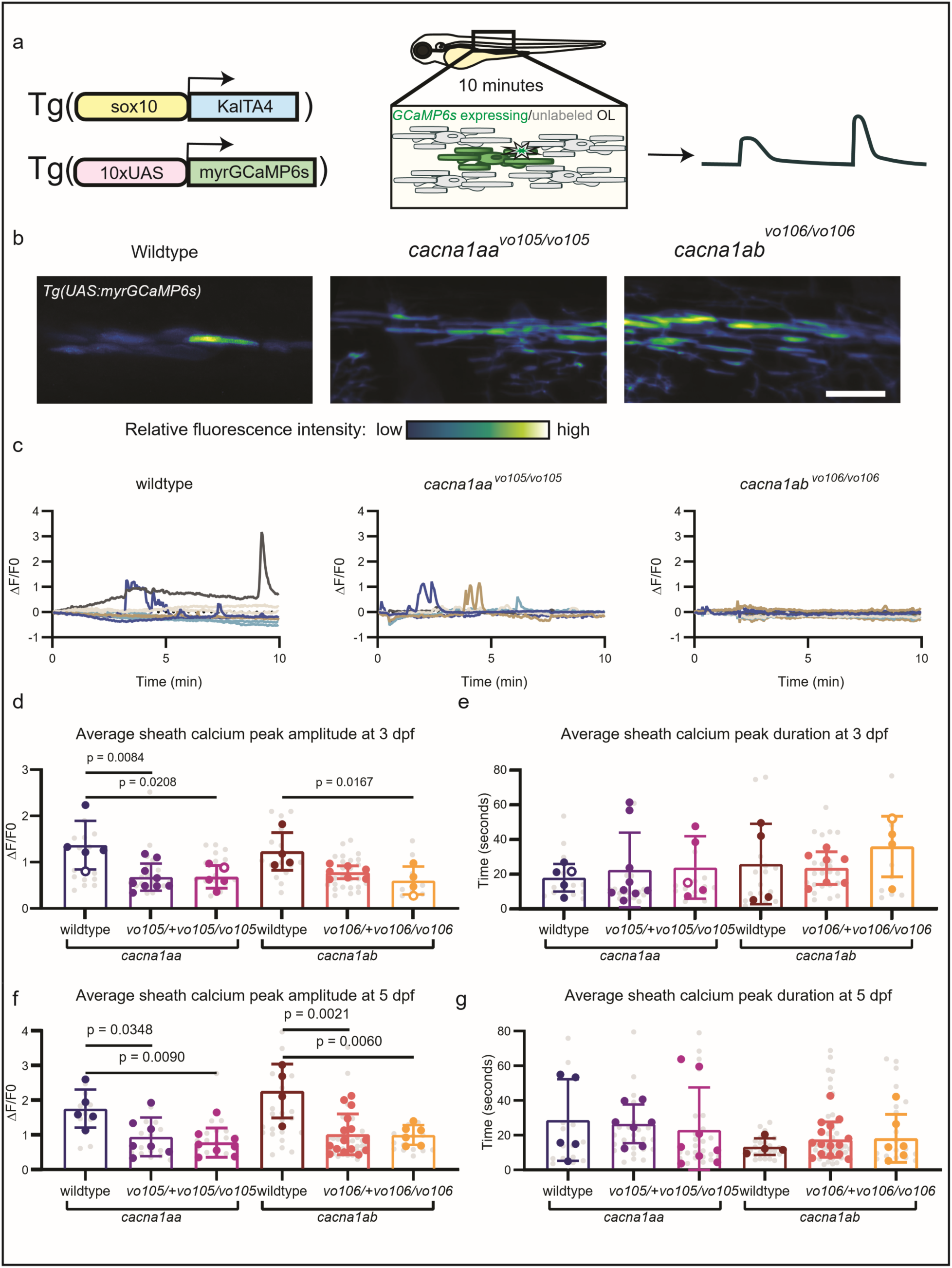
Global P/Q-type channel mutation results in reduced amplitude OL sheath calcium events. a. Schematic of experimental setup. b. Example time projections from sheath calcium imaging at 3 dpf with *Tg(sox10:KalTA4;UAS:myrGCaMP6s)*, pseudocolored by relative fluorescence intensity. c. Example traces of GCaMP6s fluorescence intensity over 10-minute recording in wild-type, *cacna1aa^vo105/vo105^*, and *cacna1ab^vo106/vo106^* animals at 3 dpf, with same examples chosen in b and c. d. Average amplitude of sheath calcium events at 3 dpf (*cacan1aa* ANOVA p = 0.0072, Tukey’s multiple comparisons wildtype vs *cacna1aa^vo105/+^*p = 0.0084, wildtype vs *cacna1aa^vo105/vo105^* p = 0.0208, *cacna1ab* ANOVA p = 0.0167, wildtype vs *cacna1ab^vo106/+^* p = 0.0506, wildtype vs *cacna1ab^vo106/vo106^* p = 0.0167). e. Average duration of sheath calcium peak events at 3 dpf (*cacna1aa* ANOVA p = 0.8627, *cacna1ab* ANOVA p = 0.4760). f. Average amplitude of sheath calcium events at 5 dpf (*cacna1aa* ANOVA p = 0.0096, Tukey’s multiple comparisons wildtype vs *cacna1aa^vo105/+^* p = 0.0348, wildtype vs *cacna1aa^vo105/vo105^* p = 0.0090, *cacna1ab* ANOVA p = 0.0021, wildtype vs *cacna1ab^vo106/+^* p = 0.0021, wildtype vs *cacna1ab^vo106/vo106^* p = 0.0060). g. Average duration of sheath calcium peak events at 5 dpf (*cacna1aa* ANOVA p = 0.8821, *cacna1ab* ANOVA p = 0.7540). For all graphs, gray points represent individual values, and colored/outlined points represent average per animal. Unfilled points represent examples shown in b and c. Statistics computed based on per-animal averages. Sample sizes are as follows. 3 dpf: *cacna1aa* wildtype N = 5 fish, *cacna1aa^vo105/+^*N = 9 fish, *cacna1aa^vo105/vo105^* N = 5 fish, *cacna1ab* wildtype N = 4 fish, *cacna1ab^vo106/+^*N = 7 fish, *cacna1ab^vo106/106^* N = 4 fish. 5 dpf: *cacna1aa* wildtype N = 5 fish, *cacna1aa^vo105/+^*N = 7 fish, *cacna1aa^vo105/vo105^* N = 8 fish, *cacna1ab* wildtype N = 4 fish, *cacna1ab^vo106/+^*N = 15 fish, *cacna1ab^vo106/106^* N = 6 fish.

In addition to testing the contribution of P/Q-type channels to myelin sheath calcium transient activity, we also measured how these channels contribute to OPC microdomain calcium signaling. Previous work has shown OPC calcium microdomain activity hotspots are associated with future myelin formation in those regions^37^. We used a previously published transgenic line, *Tg(olig1:myrGCaMP6s)* to drive expression of membrane-tethered GCaMP6s in OPCs in *cacna1aa^vo105/+^* and *cacna1ab^vo106/+^* heterozygous intercross offspring. We then recorded GCaMP6s fluorescence in spinal cord OPCs over ten minutes in larval zebrafish at 3 and 5 dpf and measured microdomain fluorescence intensity in OPC processes (Fig. 4a, b, Supplementary Video 2). At 3 dpf, the OPC calcium peak events were similar in amplitude (Fig. 4c, *cacna1aa* ANOVA p = 0.0535, *cacna1ab* ANOVA p = 0.8486) but shorter in duration in the *cacna1aa^vo105/vo105^*, *cacna1ab^vo106/+^*, and *cacna1ab^vo106/vo106^* OPCs than in the sibling wild-type OPCs (Fig. 4d, *cacna1aa* ANOVA p =0.0288, *cacna1ab* ANOVA p = 0.0251). At 5 dpf there was no difference in peak amplitude or duration (Fig. 4e, *cacna1aa* ANOVA p = 0.3002, p = 0.4687, Fig. 4f, *cacna1aa* ANOVA p = 0.8668, *cacna1ab* ANOVA p = 0.3594). The frequency of calcium peak events was similar across genotypes at both 3 and 5 dpf (Supplementary Fig. 4c, *cacna1aa* ANOVA p = 0.8820, *cacna1ab* ANOVA p = 0.6869, Supplementary Fig. 4d, *cacna1aa* ANOVA p = 0.3996, *cacna1ab* ANOVA p = 0.6640). These results suggest a role for P/Q-type channels in modulating the duration of calcium influx into OPC microdomains at 3 dpf, but not at 5 dpf.

**Figure 4.**
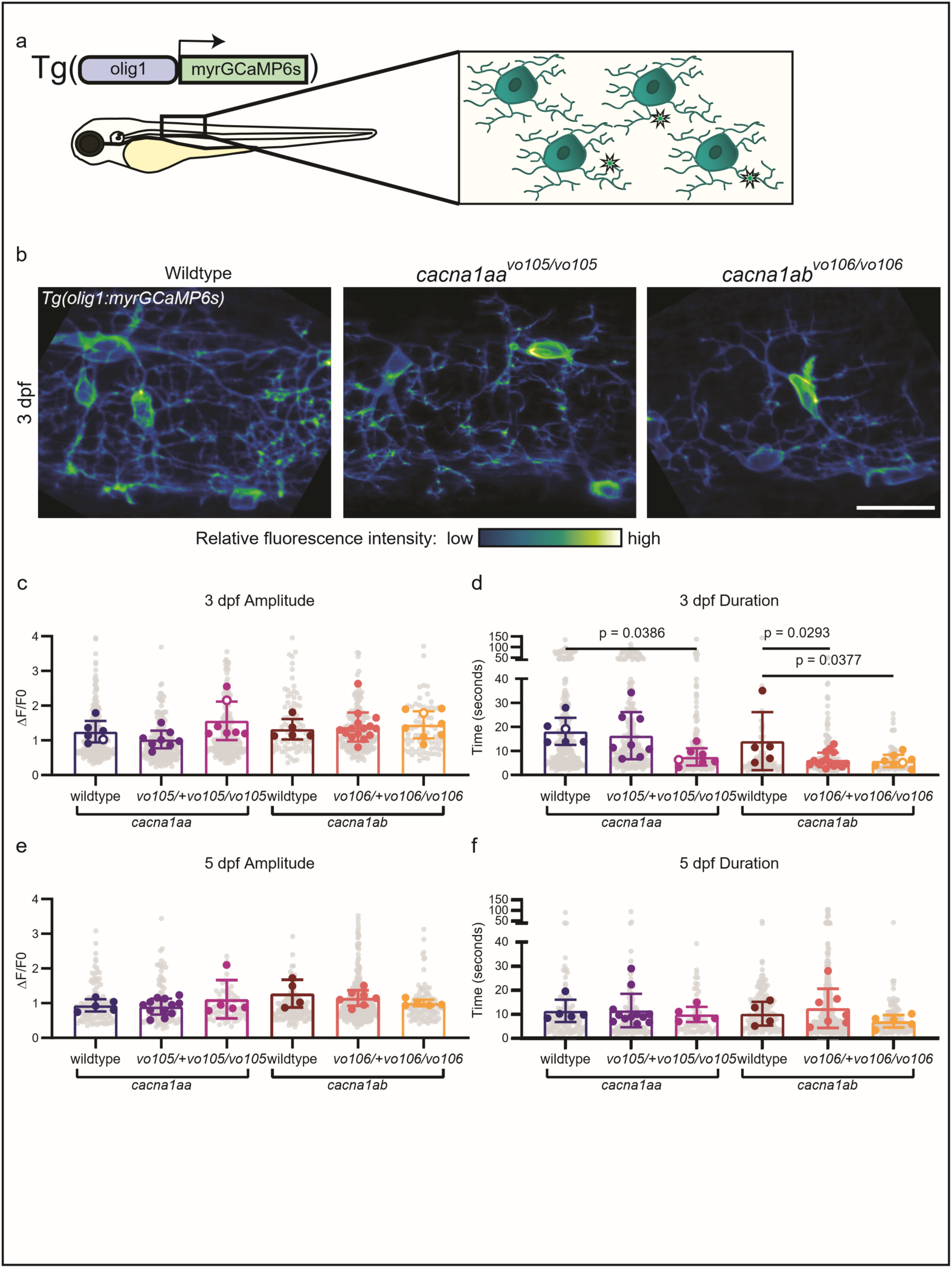
P/Q-type calcium channels modulate the duration of calcium transient events in microdomains at 3 dpf, but not 5 dpf. a. Schematic of experimental setup. b. Example time-projections from 10-minute recordings with *Tg(olig1:myrGCaMP6s)* in spinal cord of wild-type and mutant zebrafish at 3 dpf, pseudocolored. c. Amplitude of calcium microdomain peak events at 3 dpf (*cacna1aa* ANOVA p = 0.0535, *cacna1ab* ANOVA p = 0.8486). d. Duration of calcium microdomain peak events at 3 dpf (*cacna1aa* ANOVA p = 0.0288, Tukey’s multiple comparisons wildtype vs *cacna1aa^vo105/+^* p = 0.8872, wildtype vs *cacna1aa^vo105/vo105^* p = 0.0386, *cacna1ab* ANOVA p = 0.0251, wildtype vs *cacna1ab^vo106/+^* p = 0.0293, wildtype vs *cacna1ab^vo106/vo106^* p = 0.0377). e. Amplitude of 5 dpf calcium microdomain peak events (*cacna1aa* ANOVA p = 0.4687, *cacna1ab* ANOVA p = 0.3002). f. Duration of 5 dpf calcium microdomain peak events (*cacna1aa* ANOVA p = 0.8668, *cacna1ab* ANOVA p = 0.3594). For all graphs, gray points represent individual peak values, and colored/outlined points represent average per animal. Unfilled points represent examples shown in b. Statistics computed based on per-animal averages. Sample sizes are as follows. 3 dpf: *cacna1aa* wildtype N = 6 fish, *cacna1aa^vo105/+^* N = 8 fish, *cacna1aa^vo105/vo105^* N = 7 fish, *cacna1ab* wildtype N = 5 fish, *cacna1ab^vo106/+^* N = 15 fish, *cacna1ab^vo106/106^* N = 8 fish. 5 dpf: *cacna1aa* wildtype N = 5 fish, *cacna1aa^vo105/+^* N = 12 fish, *cacna1aa^vo105/vo105^* N = 5 fish, *cacna1ab* wildtype N = 4 fish, *cacna1ab^vo106/+^* N = 7 fish, *cacna1ab^vo106/106^* N = 5 fish.

In addition to transducing voltage changes into biologically important intracellular Ca^2+^ transients, VGCC membrane currents may also shape the membrane potential directly. To further investigate the membrane signaling properties of the P/Q-type channel mutant OPCs, we performed whole-cell patch-clamp electrophysiology in mutant and wild-type spinal cord OPCs at 3 dpf (Fig. 5a-c). The vast majority of OPC calcium transient activity is found in the processes, not the cell soma^37^ and VGCCs are less abundant than other ion channels in OPCs^49,50^. We did not observe whole-cell VGCC currents in wild-type or mutant OPCs. Moreover, voltage-sensitive currents were similar between wild-type and heterozygous or mutant OPCs (Fig. 5d-g). These results suggest P/Q-type channel mutation does not significantly affect the basic bioelectrical properties of these cells.

**Figure 5.**
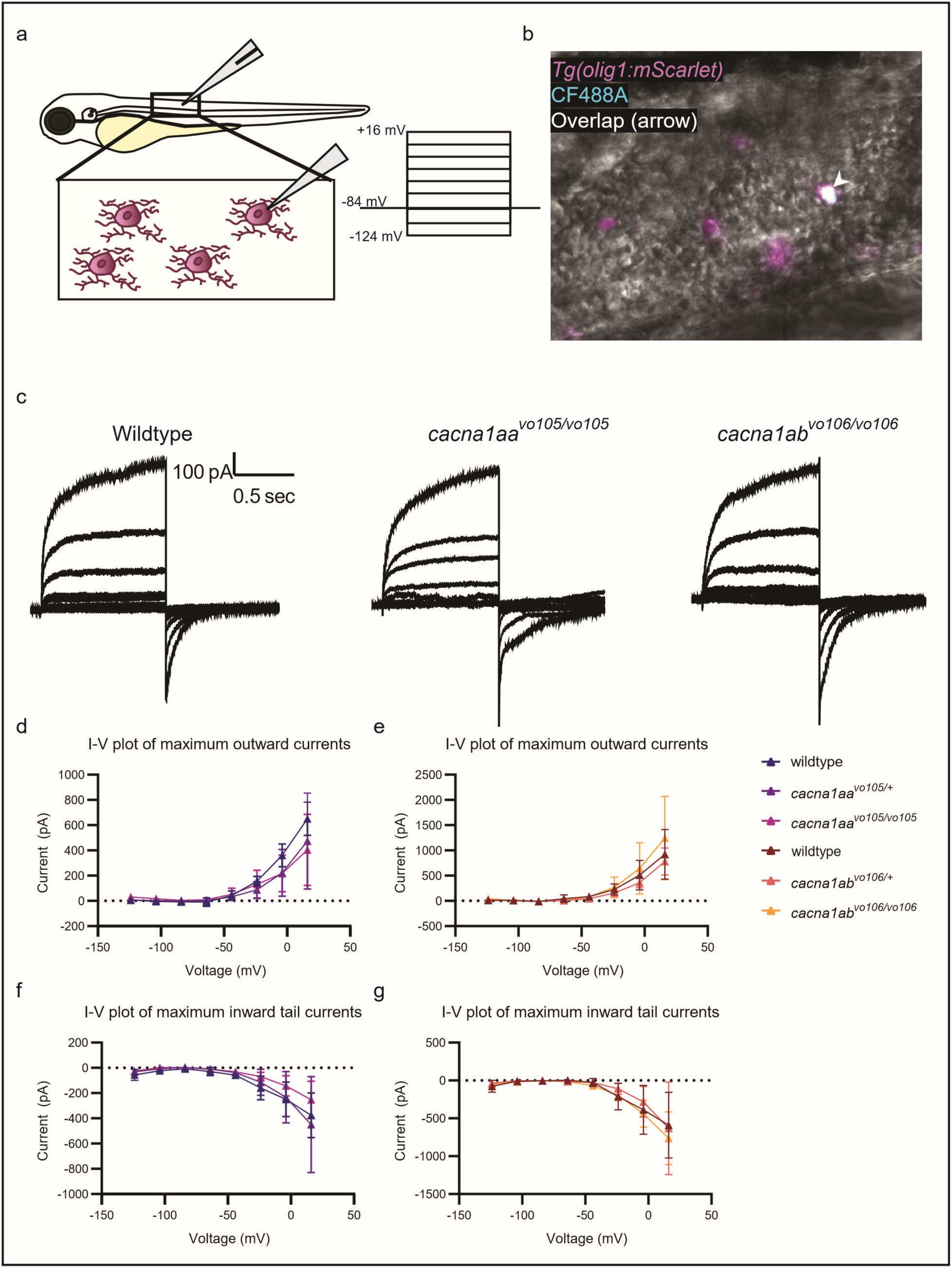
Bioelectrical properties of OPCs are similar between wild-type and mutant cells in the zebrafish spinal cord at 3 dpf. a. Schematic of experimental setup. b. Example image of patched OPC labeled with *Tg(olig1:mScarlet)* with dye fill (CF488) following recording. c. Example leak subtracted (P/4) currents from voltage clamp recordings of OPCs according to voltage step schematic in a. d. I-V plot of maximum outward currents from voltage clamp recordings across *cacna1aa* genotypes (ANOVA p = 0.5703). e. I-V plot of maximum outward currents from voltage clamp recordings across *cacna1ab* genotypes (ANOVA p = 0.4192). f. I-V plot of maximum inward tail currents recorded in voltage clamp across *cacna1aa* genotypes (ANOVA p = 0.5514). g. I-V plot of maximum inward tail currents recorded in voltage clamp across *cacna1ab* genotypes (ANOVA p = 0.7623). Sample sizes are as follows, with one cell per animal: *cacna1aa* wildtype N = 4 fish, *cacna1aa^vo105/+^*N = 3 fish, *cacna1aa^vo105/vo105^* N = 3 fish, *cacna1ab* wildtype N = 4 fish, *cacna1ab^vo106/+^*N = 6 fish, *cacna1ab^vo106/106^* N = 3 fish.

Similarly, P/Q-type channel mutations did not change the number of OLCs at 3 or 5 dpf (Supplementary Fig. 5a, b *cacna1aa* ANOVA p = 0.5276, *cacna1ab* ANOVA p = 0.7847, Fig. 3c *cacna1aa* ANOVA p = 0.6958, *cacna1ab* ANOVA p = 0.9587). There was also a similar number of OPCs at 3 dpf in wild-type and mutant spinal cords (Supplementary Fig 5d, e, *cacna1aa* ANOVA p = 0.3279, *cacna1ab* ANOVA p = 0.1413). These results suggest the proliferation and survival of OLCs is not grossly abnormal in the P/Q-type channel mutants. The OLC calcium signaling differences between mutant and wild-type OLCs appears to affect the development of myelin sheaths, but not to have a great effect on the membrane properties, proliferation, or survival of these cells. We next sought to understand the ways myelin sheath development differs between P/Q-type channel mutant and wild-type OLs.

### Early myelinating OLs require P/Q-type channels for normal sheath elongation

Because the *cacna1aa^vo105^* and *cacna1ab^vo106^* heterozygous and homozygous mutant OLs exhibit deficits in myelin production, we next tested how these mutations affect sheath elongation in newly myelinating OLs. We hypothesized that disruption of calcium signaling in the differentiating OL may impair actin cytoskeletal organization and initial ensheathment of axons. To test this idea, we performed time-lapse *in vivo* imaging for eight hours overnight from 2-3 dpf in *cacna1aa^vo105/+^* and *cacna1ab^vo106/+^* heterozygous intercross offspring zebrafish injected with a construct to sparsely label OLC membrane and F-actin (*sox10:myrEGFP-P2A-Lifeact-TagRFP*) (Fig. 6a, b, Supplementary Video 3). We found that *cacna1aa^vo105/vo105^*, *cacna1ab^vo106/+^*, and *cacna1ab^vo106/vo106^* OLs exhibited a slower net rate of sheath elongation during the imaging period than corresponding wild-type controls (Fig. 6c, *cacna1aa* ANOVA p = 0.0074, *cacna1ab* ANOVA p <0.0001). This reduced ensheathment length is likely due to a failure to stabilize and elongate newly formed sheaths, as the number of retracted ensheathments and new ensheathments formed during imaging were similar across genotypes (Fig. 6d, *cacna1aa* ANOVA p = 0.8657, *cacna1ab* ANOVA p = 0.2019, Fig. 6e, *cacna1aa* ANOVA p = 0.2822, *cacna1ab* ANOVA p = 0.8473). The actin cytoskeleton as visualized by Lifeact-TagRFP moved to a similar degree across genotypes (Fig. 6f, *cacna1aa* ANOVA p = 0.1626, *cacna1ab* ANOVA p = 0.8903), suggesting the wrapping dynamics of these newly formed sheaths were not significantly affected by P/Q-type channel mutation. These results indicate that the disrupted calcium signaling in the *cacna1aa^vo105/+^*, *cacna1aa^vo105/vo105^*, *cacna1ab^vo106/+^*, and *cacna1ab^vo106/vo106^* OLCs (Fig. 3, 4) likely negatively affects calcium-dependent sheath stabilization that facilitates the elongation of new sheaths.

**Figure 6.**
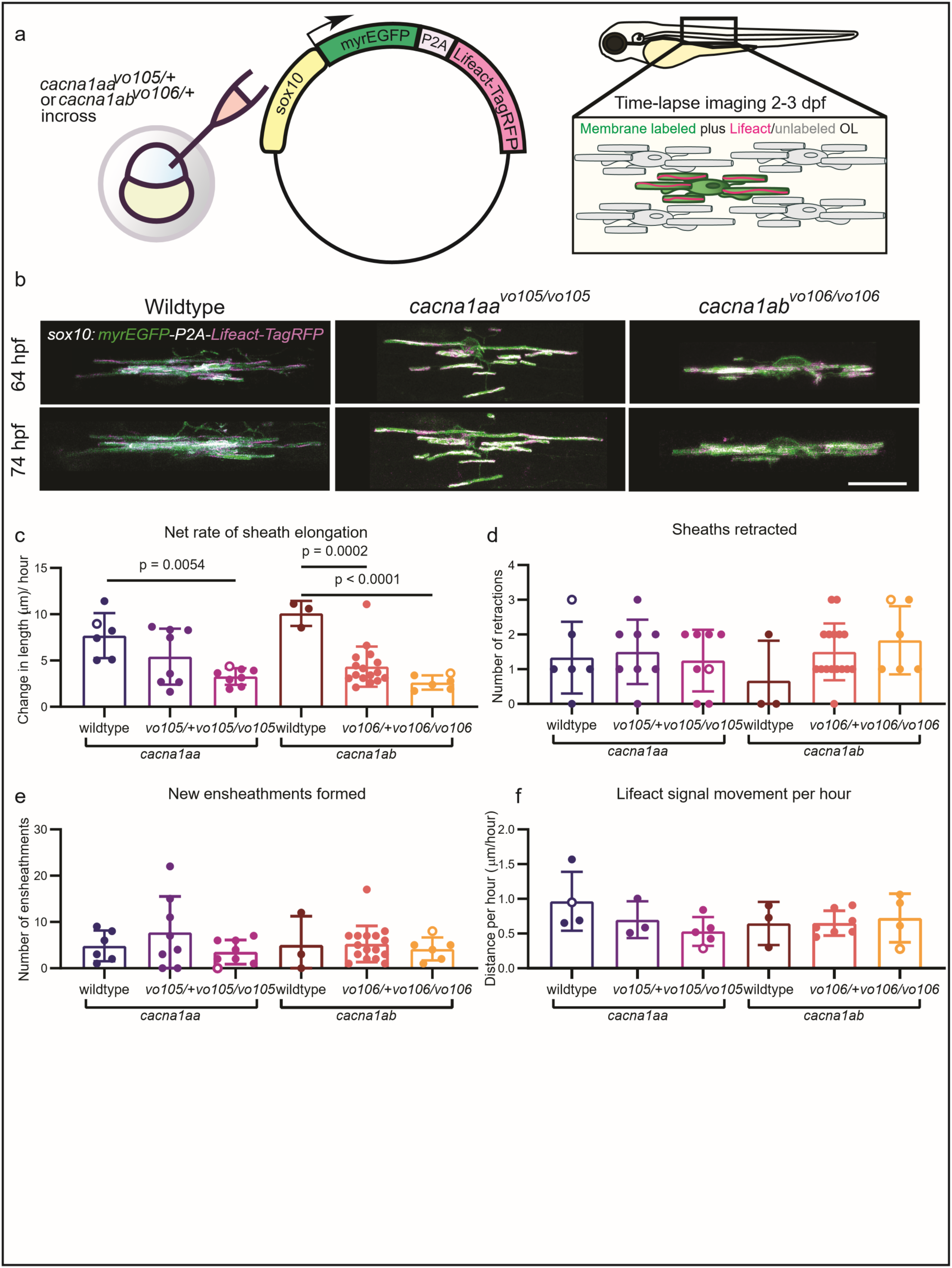
P/Q-type channel mutants exhibit reduced myelin sheath elongation in newly differentiated OLs. a. Schematic of experimental setup. b. Example images of OLs taken from 2-3 dpf time-lapse imaging. c. Net rate of sheath elongation in µm per hour during imaging period (*cacna1aa* ANOVA p = 0.0074, Tukey’s multiple comparisons wildtype vs *cacna1aa^vo105/vo105^* p = 0.0054, *cacna1ab* ANOVA p < 0.0001, wildtype vs *cacna1ab^vo106/+^*p = 0.0002, wildtype vs *cacna1ab^vo106/vo106^* p < 0.0001). d. Number of sheaths retracted by each cell (*cacna1aa* ANOVA p = 0.8657, *cacna1ab* ANOVA p = 0.2019). e. Number of new sheaths formed during imaging (*cacna1aa* ANOVA p = 0.2822, *cacna1ab* ANOVA p = 0.8473). f. Lifeact signal movement distance per hour relative to bottom of sheath, averaged along the length of the sheath (*cacna1aa* ANOVA p = 0.1626, *cacna1ab* ANOVA p = 0.8903). Each data point represents per-cell average and unfilled points represent examples shown in b. For all graphs except f, Sample size is as follows: *cacna1aa* wildtype N = 5 fish, 11 cells, *cacna1aa^vo105/+^*N = 6 fish, 11 cells, *cacna1aa^vo105/vo105^* N = 4 fish, 10 cells, *cacna1ab* wildtype N = 2 fish, 3 cells, *cacna1ab^vo106/+^*N = 8 fish, 16 cells, *cacna1ab^vo106/106^* N = 3 fish, 5 cells. For f, sample size is as follows: *cacna1aa* wildtype N = 2 fish, 4 cells, *cacna1aa^vo105/+^*N = 3 fish, 4 cells, *cacna1aa^vo105/vo105^* N = 4 fish, 5 cells, *cacna1ab* wildtype N = 2 fish, 3 cells, *cacna1ab^vo106/+^* N = 4 fish, 5 cells, *cacna1ab^vo106/106^* N = 3 fish, 4 cells.

### P/Q-type channel mutants have increased OL membrane blebbing associated with sheath shortening

Next, we compared remodeling of myelin sheaths in *cacna1aa^vo105^* and *cacna1ab^vo106^* OLs using fish with the transgenic *Tg(mbp:EGFP-CAAX)* background to label OL membrane (Fig. 7a). We imaged the spinal cords of these larvae at 5 dpf and observed membrane blebbing associated with myelin sheaths (Fig. 7b). This membrane blebbing resembles previously reported membrane outfoldings that occur at a low frequency during normal development^51^, but the dynamics of these structures are not well understood. We performed time-lapse imaging *in vivo* from 5-6 dpf (Fig. 7a, c, Supplementary Video 4), and found that in all cases, sheaths with membrane blebbing shortened over the course of the eight-hour time-lapse (Fig. 7d, e, ANOVA p < 0.0001). These results suggest that this membrane blebbing is a hallmark of rapid sheath shortening, and their increased incidence in P/Q-type channel mutants could contribute to the shorter sheath lengths seen in P/Q-type channel mutant OLs.

**Figure 7.**
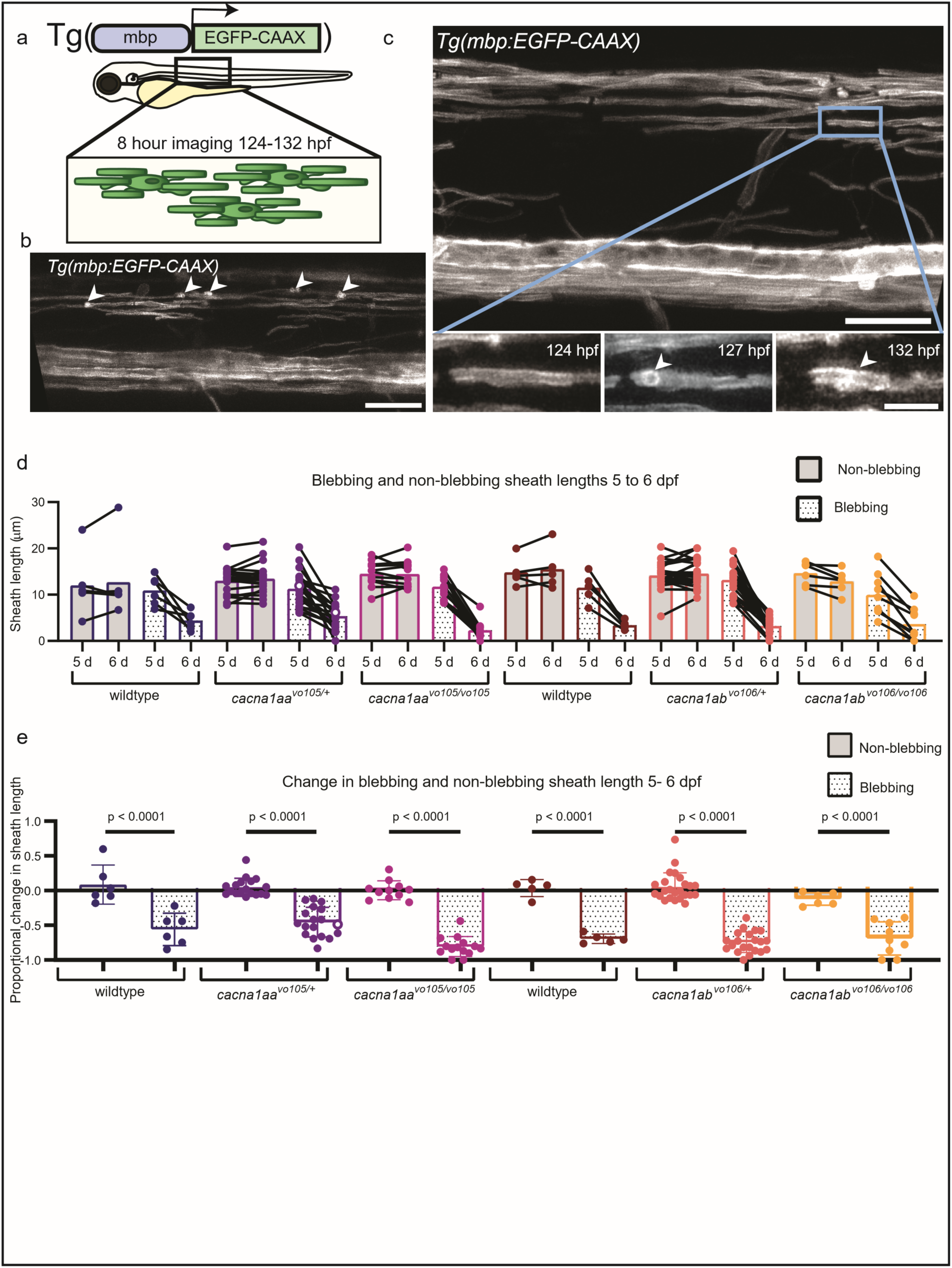
P/Q-type channel mutant OLs exhibit membrane blebbing associated with sheath shortening. a. Schematic of experimental setup. b. Example image of membrane blebbing structures from a *Tg(mbp:EGFP-CAAX)* transgenic, *cacna1ab* crispant (F0) spinal cord at 5 dpf, arrowheads labeling blebbing membrane. c-e. Analysis in stable mutants. c. Example of membrane blebbing appearing over the course of an 8-hour time-lapse between 5-6 dpf. d. Comparison of sheath lengths at the beginning and end of time-lapse imaging in sheaths associated with a blebbing and randomly selected non-blebbing control sheaths from the same fish. e. Fold change of sheath lengths over the course of the time-lapse in blebbing sheaths and non-blebbing controls (ANOVA p < 0.0001, Tukey’s multiple comparisons p < 0.0001 for each). Each point or pair on graph represents one myelin sheath. Unfilled points represent example shown in c. Sample sizes are as follows: *cacna1aa* wildtype N = 3 fish, 6 blebbing/non-blebbing sheath pairs, *cacna1aa^vo105/+^* N = 10 fish, 21 blebbing/20 non-blebbing sheath pairs, *cacna1aa^vo105/vo105^* N = 5 fish, 14 blebbing/11 non-blebbing sheath pairs, *cacna1ab* wildtype N = 2 fish, 5 blebbing/non-blebbing sheath pairs, *cacna1ab^vo106/+^* N = 12 fish 21 blebbing/ 24 non-blebbing sheaths, *cacna1ab^vo106/106^* N = 3 fish, 9 blebbing/6 non-blebbing sheath pairs.

We observed membrane blebbing occurring at a low frequency in the wild-type OLs, but they occurred more frequently in *cacna1aa^vo105/vo105^* and *cacna1ab^vo106/vo106^* OLs than in wildtype (Fig 8a-c, *cacna1aa* ANOVA p = 0.0262, *cacna1ab* ANOVA p = 0.0371). This result suggests that this may be a normal developmental phenomenon that is exaggerated in P/Q-type channel mutants. The membrane blebbing occurs closer to the edge of the sheath than would occur by random chance (Fig. 8d, one-sample t-test p < 0.0001), which may indicate a relationship between disorganization of the membrane and perinodal organization in these developing sheaths. This membrane blebbing was also more prevalent in cell-specific mutated cells than un-mutated control (Fig. 8e, f, ANOVA p < 0.0001), which suggests a cell-autonomous regulation of OL membrane blebbing.

**Figure 8.**
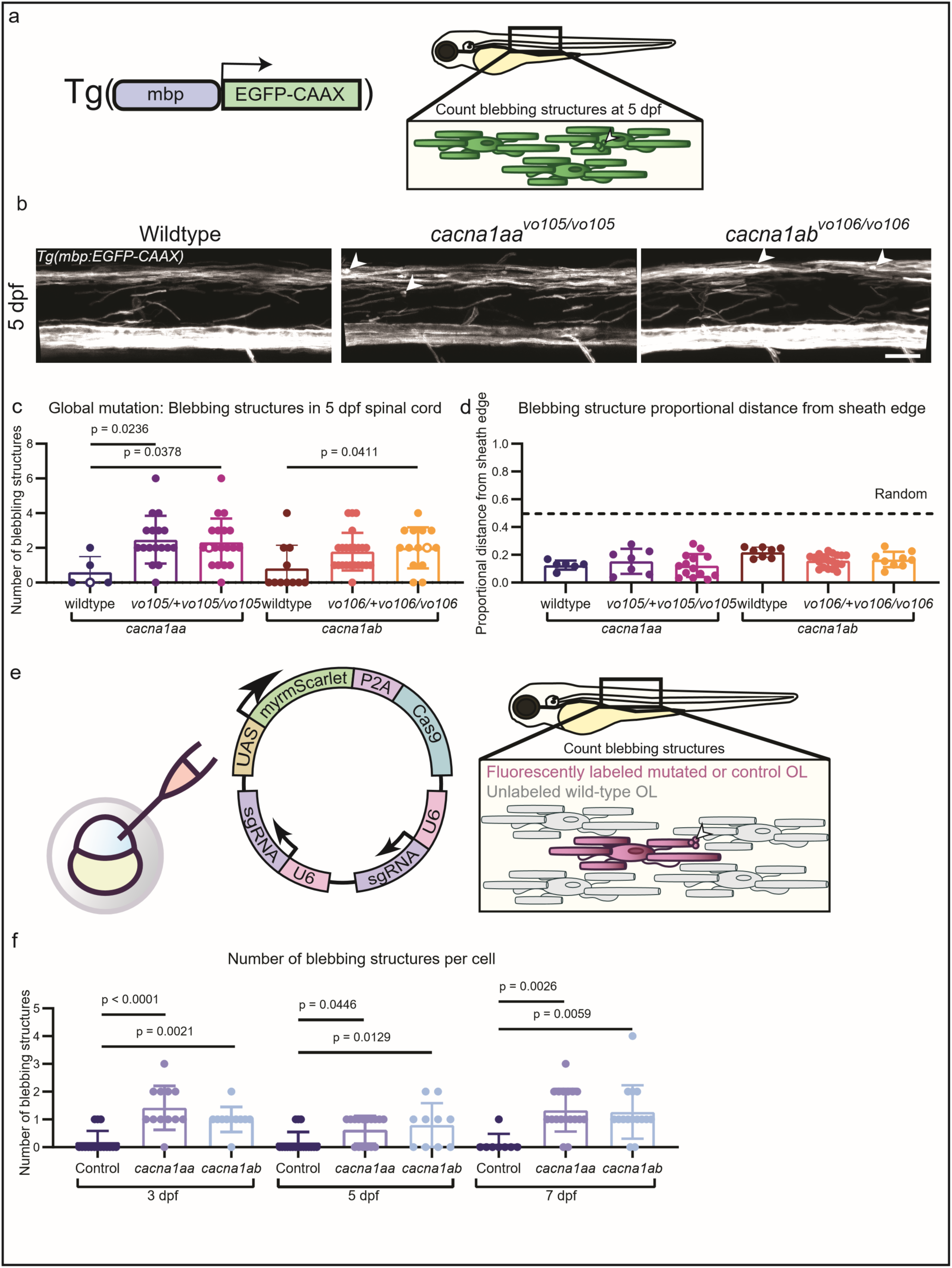
Myelin-associated membrane blebbing occurs more frequently in P/Q-type channel mutant OLs than wild-type OLs. a. Schematic of experimental setup for global mutant transgenic imaging of membrane blebbing at 5 dpf. b. Example images of *Tg(mbp:EGFP-CAAX)* transgenic fish at 5 dpf with wild-type or stable global mutant backgrounds, with membrane blebbing (arrowheads) in homozygous mutant animals. c. Quantification of membrane blebbing in *Tg(mbp:EGFP-CAAX)* transgenic fish at 5 dpf in one somite across genotypes (*cacna1aa* ANOVA p = 0.0262, Tukey’s multiple comparisons wildtype vs *cacna1aa^vo105/+^*p = 0.0236, wildtype vs *cacna1aa^vo106/vo106^* p = 0.0378, *cacna1ab* ANOVA p = 0.0371, wildtype vs *cacna1ab^vo105/+^* p = 0.0738, wildtype vs *cacna1ab^vo106/vo106^* p = 0.0411). Sample sizes are as follows: *cacna1aa* wildtype N = 5 fish, *cacna1aa^vo105/+^*N = 17 fish, *cacna1aa^vo105/vo105^* N = 19 fish, *cacna1ab* wildtype N = 11 fish, *cacna1ab^vo106/+^*N = 18 fish, *cacna1ab^vo106/106^* N = 11 fish. d. Quantification of distance from blebbing to edge of sheath at time blebbing first appeared, compared with random (average across genotypes compared with hypothetical random mean = 0.5, one-sample t-test p < 0.0001). Sample size as follows: *cacna1aa* wildtype N = 3 fish, 6 sheaths, *cacna1aa^vo105/+^* N = 3 fish, 7 sheaths, *cacna1aa^vo105/vo105^* N = 5 fish, 14 sheaths, *cacna1ab* wildtype N = 4 fish, 8 sheaths, *cacna1ab^vo106/+^* N = 7 fish, 12 sheaths, *cacna1ab^vo106/106^* N = 3 fish 9 sheaths. e. Experimental setup for cell-specific mutants to quantify membrane blebbing. f. Quantification of blebbing structures in cell-specific mutants at 3, 5, and 7 dpf (3 dpf ANOVA p < 0.0001, Tukey’s multiple comparisons control vs *cacna1aa* p < 0.0001, control vs *cacna1ab* p = 0.0021, 5 dpf ANOVA p = 0.0082, control vs *cacna1aa* p = 0.0446, control vs *cacna1ab* p = 0.0129, 7 dpf ANOVA p = 0.0023, control vs *cacna1aa* p = 0.0026, control vs *cacna1ab* p = 0.0059). For c, each point on graph represents one animal. Unfilled points represent examples shown in b. For d, each point on graph represents one myelin sheath. For f, each point on graph represents one cell.

Together, these results indicate that P/Q-type channels play an important role in regulating developmental myelination. They regulate the calcium signaling in developing myelin sheaths, as well as modulate the lengthening of new sheaths and the incidence of membrane blebbing associated with sheath shortening. Further study into this dynamic blebbing, its relationship to calcium signaling, and its role in normal and pathological myelin development is warranted.

## Discussion

OLC calcium signaling is a context-dependent, complex signaling process that impacts a multitude of cellular functions ^6,8,11,22,48,52^. Here, we show that P/Q-type voltage-gated calcium channels regulate multiple aspects of developmental myelination. Signaling via P/Q-type channels positively affects the amplitude of OL myelin sheath calcium events and extends the duration of OPC calcium microdomain transients. These effects on OLC calcium signaling fit with a previously proposed model of voltage-gated calcium channel function in OLCs wherein activation of calcium-permeable AMPA receptors caused a local depolarization sufficient to activate voltage-gated calcium channels^21^. In this model, most calcium influx occurs via the AMPA receptor, but the voltage gated calcium channels activated downstream could modulate the amplitude and/or duration of the signal, as we see in our results. In addition to calcium imaging with genetically encoded calcium indicators, we used whole-cell patch-clamp electrophysiology of zebrafish OPCs *in vivo*, which, to our knowledge, represents the first use of this technique in these cells in zebrafish. Our results show the basic bioelectrical properties of OPCs were similar between wild-type and P/Q-type channel mutant animals, which is not surprising considering the low abundance of these channels compared to other channels in OPCs. These results also provide evidence that OPCs in the zebrafish spinal cord at 3 dpf have similar bioelectrical properties to analogous timepoints in mammals, including similar I-V curve trends as those reported in rodent OPCs at early postnatal timepoints, as has been previously reported^49,50^. Beyond P/Q-type calcium channels, this technique could also provide further insight into the significance of calcium transients and more in future studies. These results suggest that the local calcium transient signaling changes to OLCs likely play a larger role in modulating the myelination effects, rather than larger changes to the membrane properties of these cells.

Several questions remain, including what is downstream of changes in calcium signaling mediated by P/Q-type channels. Previous studies have shown there are many processes regulated by calcium that play vital roles in myelin development, including vesicle cycling, gene transcription, and cytoskeletal organization^16–20^. It is possible that each of these processes could regulate the changes in myelin development we describe here. Exocytosis has been described as an important process in the rapid membrane expansion required for myelin formation^3,4^, and we know this process can be calcium-dependent^54,55^. The onset of myelination involves extensive gene transcription and translation to produce myelin basic protein and many other important components of myelin^56^. Energetic support for myelination via modulation of mitochondria can also be a calcium-dependent process^57^. Cytoskeletal organization is likewise a vital part of myelin sheath formation and organization^1,58,59^. Calcium is important for a wide range of cellular processes that regulate myelination, and based on our results and others’, it is clear that the disruption of calcium signaling leads to impaired myelination. Further study of the many ways in which calcium signaling occurs in OLCs will give a better understanding of the significance of calcium activity to specific downstream cell processes, as well as the types of signals and contexts that work in concert to regulate myelination.

Myelin integrity is important for the proper function of the vertebrate nervous system, and there are a broad array of pathologies in which myelination is disrupted. The absence of myelination can cause neuronal damage and dysfunction, either through a demyelinating disorder like multiple sclerosis or due to deficits in developmental myelination like leukodystrophies. Abnormal myelin formation can also cause problems, for example through maladaptive myelination in epilepsy. Myelin sheath length is an important property of myelin that affects the efficiency of neuronal signal transmission by altering the distance between nodes of Ranvier and can have effects on signal coordination^9,60^.

Our findings show P/Q-type channel mutations reduce average sheath length in developing OL myelin via a cell-autonomous mechanism and reduced total myelin output in OLs in both lineage-specific and global mutant approaches. By electron microscopy, we also found there are fewer ventral axons myelinated in the *cacna1aa^vo105/vo105^* and *cacna1ab^vo106/106^* animals compared to wild-type siblings. Together, these results demonstrate the importance of P/Q-type channels in OL development and myelin formation. In our results, we found two effects of P/Q-type channel mutation on myelin development that likely contribute to these changes in *cacna1aa^vo105^* and *cacna1ab^vo106^* heterozygous and homozygous mutant spinal cord: reduced rate of myelin formation during early myelination, and increased incidence of membrane blebbing associated with myelin sheath shortening. In the case of reduced rate of myelin formation, other studies have shown that calcium signaling is required in early sheath development to stabilize new sheaths and promote sheath elongation^5,6^. Our results fit into this established paradigm, with a decrease in sheath calcium peak amplitude correlating with a reduced stabilization and elongation of new sheaths. Further research into the fine-tuning of these processes is needed to understand the mechanisms underlying tight control of sheath dynamics by calcium signals.

The other major myelin deficit we describe in the *cacna1aa^vo105^* and *cacna1ab^vo106^* heterozygous and homozygous mutant spinal cord is blebbing, an OL membrane phenomenon associated with rapid myelin sheath shortening during development. This process appears to be somewhat physiological, in that it is seen under normal conditions, but increases significantly in the *cacna1aa^vo105^* and *cacna1ab^vo106^* heterozygous and homozygous mutant spinal cord. We suspect this membrane blebbing likely represents membrane outfoldings, as it appears similar to previously described disorganized myelin structures^51,58^. To our knowledge, this OL membrane blebbing has not previously been described in association with sheath shortening, though in our study this association is robust. It is possible that the rapid sheath shortening seen in our experiments causes the membrane to collapse into these blebbing structures. There is growing evidence in the field that disruption to ion homeostasis, e.g. by mutating ion channels, can have a great impact on myelination^48,61,62^. Further study to understand the significance of this phenomenon in myelin development, and in the overall function of the nervous system will help determine if this membrane blebbing represents pathological adaptations in the absence of normal OL calcium signaling.

Interestingly, the phenotypes we characterize here are similar between heterozygous and homozygous mutant *cacna1aa^vo105^* and *cacna1ab^vo106^* animals in many cases (e.g., Fig. 1c, e, f, g, Fig. 3d, f, Fig. 8c). Since many CACNA1A-related disorders result from haploinsufficiency due to loss of function mutations, it makes sense that a heterozygous mutation could cause significant developmental changes. Because the heterozygous *cacna1aa^vo105/+^*and *cacna1ab^vo106/+^* animals displayed intermediate phenotypes in some cases (e.g., Fig. 1h, Fig. 6c), the *cacna1aa^vo105^* and *cacna1ab^vo106^* mutations in this study likely function by reducing gene dosage, where the threshold for genetic perturbation inducing developmental OLC changes is relatively low.

Our results also raise the possibility of OLC and/or myelin contribution to the pathogenesis of CACNA1A-related disorders, which has not previously been examined to our knowledge. Further study of the myelin changes in context of motor activity, seizure susceptibility, and other effects in animal models could provide useful insight into the pathogenesis of CACNA1A-related disorders. Given previous studies showing a role for OLCs in reinforcing seizure networks through maladaptive myelination^63^, it is possible the deficits in myelination described here could play a role in pathogenesis of CACNA1A-related disorders. If OLCs are indeed involved in this process, future work could uncover new possible therapeutic targets.

## Acknowledgements

We are grateful to past and current members of the Monk lab, including Suhail Akram, Emma Brennan, Austin Forbes, Kyla Hamling, Tia Perry, and Adriana Reyes for zebrafish care. We would like to thank the Advanced Light Microscopy Core at OHSU (RRID: SCR_009961). This project was supported by the National Institutes of Health/National Institute of Neurological Disorders and Stroke (F31NS137600 to M.P.; F32NS123005 to C.L.C.; R01NS120651 to K.R.M.) and by grants awarded by the U.S. Department of Veterans Affairs (BX002547 to S.M.S.).

---

<u>Supplementary Video 1.</u> Example videos of OL sheath calcium recordings with *Tg(sox10:KalTA4;UAS:myrGCaMP6s)*, as seen in Figure 3.

<u>Supplementary Video 2.</u> Example videos of OPC microdomain calcium recordings with *Tg(olig1:myrGCaMP6s)*, as seen in Figure 4.

<u>Supplementary Video 3.</u> Example video of sheath elongation over 2.5 hours in 3 dpf time-lapse with *sox10:myrEGFP-P2A-Lifeact-TagRFP,* as seen in Figure 6.

<u>Supplementary Video 4.</u> Example video of membrane blebbing in association with myelin sheath shortening using *Tg(mbp:EGFP-CAAX)* as seen in Figure 7.

**Supplementary Figure 1.**
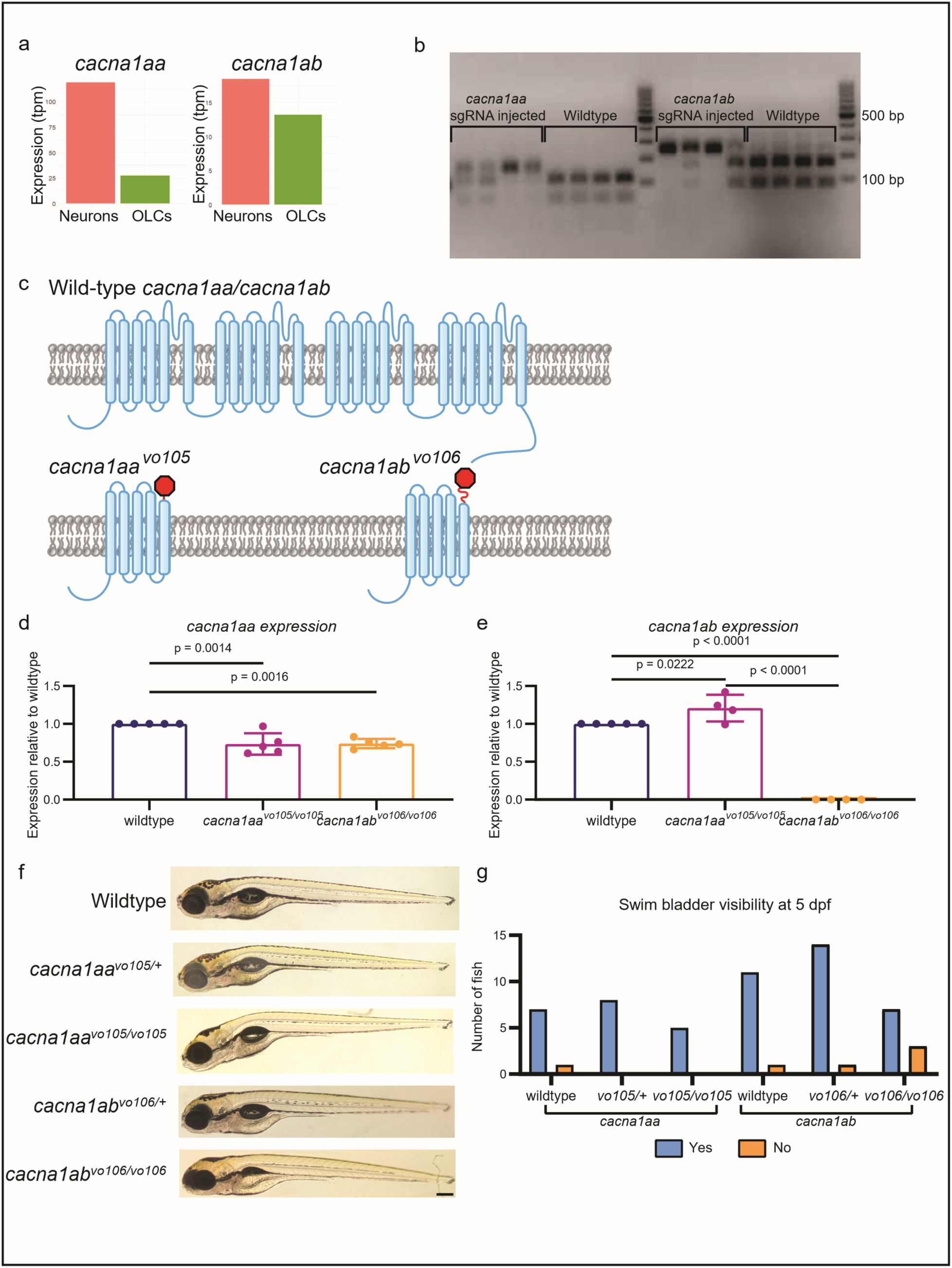
P/Q-type channel genes are expressed by OLCs, and global mutation of these genes results in reduced expression. a. Bulk transcriptomics data of P/Q-type channel gene expression in zebrafish at 3 dpf^46,47^. b. Gel demonstrating efficiency of *cacna1aa* and *cacna1ab* sgRNA used in CRISPR/Cas9 cell-specific mutagenesis system, generation of global mutants. c. Cartoon representation of predicted gene products from *cacna1aa^vo105^* and *cacna1ab^vo106^* alleles compared to wildtype. d. RT-qPCR results measuring *cacna1aa* expression in pooled 5 dpf larvae across genotypes (ANOVA p = 0.0006, Tukey’s multiple comparisons test wildtype vs *cacna1aa^vo105/vo105^* p = 0.0014, wildtype vs *cacna1ab^vo106/vo106^*p = 0.0016, *cacna1aa^vo105/vo105^* vs *cacna1ab^vo106/vo106^* p = 0.9972). e. qPCR results of *cacna1aa* and *cacna1ab* gene expression in homozygous mutants vs wild-type zebrafish embryos at 3 dpf (ANOVA p < 0.0001, Tukey’s multiple comparisons test wildtype vs *cacna1aa^vo105/vo105^* p = 0.0222, wildtype vs *cacna1ab^vo106/vo106^* p < 0.0001, *cacna1aa^vo105/vo105^* vs *cacna1ab^vo106/vo106^* p < 0.0001). Wildtype N = 16 pooled larvae, 6 technical replicates, *cacna1aa^vo105/vo105^* N = 12 pooled larvae, 6 technical replicates, *cacna1ab^vo106/vo106^* N = 6 pooled larvae, 6 technical replicates. f. Gross morphology of zebrafish larvae at 5 dpf (scale bar = 2 mm). g. Number of fish of each genotype with visible or non-visible swim bladder at 5 dpf (Chi-Square p = 0.1136. *cacan1aa*: wildtype N = 8, *vo105/+* N = 10, *vo105/vo105* N = 7; *cacna1ab*: wildtype N = 12, *vo106/+* N = 15, *vo106/vo106* N = 8).

**Supplementary Figure 2.**
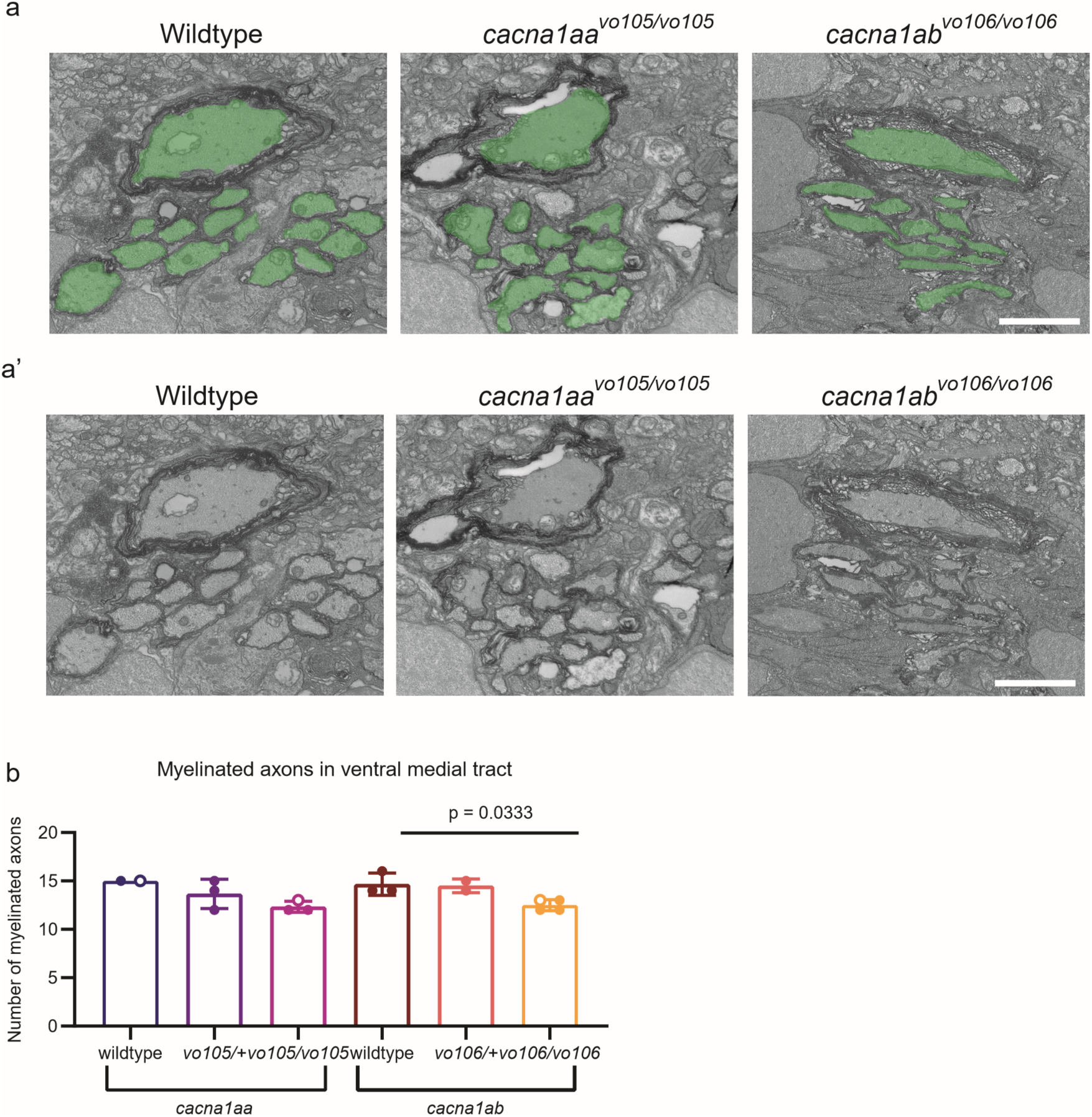
There are fewer myelinated ventral axons in P/Q-type channel mutants compared to wildtype. a. Example images of ventral spinal cord transmission electron micrographs in 5 dpf wild-type, *cacna1aa^vo105/105^*, and *cacna1ab^vo106/vo106^* zebrafish, with counted myelinated axons pseudocolored in green. a’. Images from a without pseudocoloring. b. Quantification of myelinated axons in the ventral medial tract in each genotype, *cacna1aa* ANOVA p = 0.0896; *cacna1ab* ANOVA p = 0.0264, Tukey’s multiple comparisons test wildtype vs *cacna1ab^vo106/+^* p = 0.9740, wildtype vs *cacna1ab^vo106/vo106^* p = 0.0333, *cacna1ab^vo106/+^* vs *cacna1ab^vo106/vo106^* p = 0.0724). Sample sizes for *cacna1aa:* wildtype N = 2, *vo105/+* N= 3, *vo105/vo105* N = 3; *cacna1ab:* wildtype N = 3, *vo106/+* N = 2, *vo106/vo106* N = 4.

**Supplementary Figure 3.**
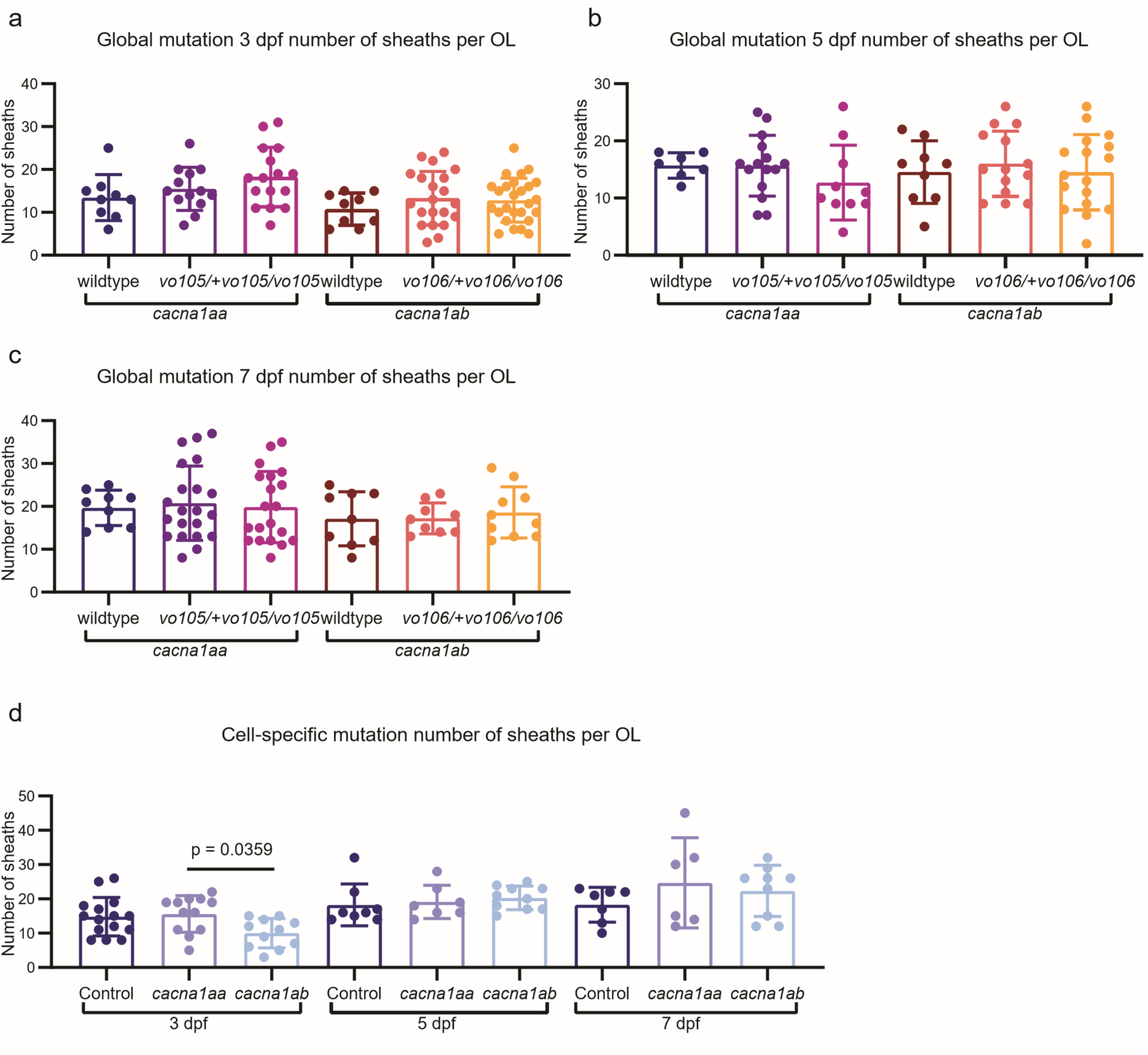
Myelin sheath numbers per OL are largely similar across genotypes in global mutation and conditions in cell-specific mutants. Comparison of number of sheaths per OL in globally mutant sparsely labeled OLs, as seen in Fig. 1, shows no significant differences across groups at a. 3 dpf (*cacna1aa* ANOVA p = 0.1524, *cacna1ab* ANOVA p = 0.5006), b. 5 dpf (*cacna1aa* ANOVA p = 0.3477, *cacna1ab* ANOVA p = 0.7691) or c. 7 dpf (*cacna1aa* ANOVA p = 0.9116, *cacna1ab* ANOVA p = 0.8020). d. Comparison of number of sheaths per OL in cell-specific mutant OLs, as seen in Fig. 2, shows a difference between *cacna1aa-*targeted and *cacna1ab-*targeted OLs at 3 dpf, but no other significant differences across groups (3 dpf ANOVA p = 0.0279, Tukey’s multiple comparisons test control vs *cacna1aa-*targeted p = 0.9189, control vs *cacna1ab-*targerted p = 0.0627, *cacna1aa-*targetred vs *cacna1ab-*targeted p = 0.0359. 5 dpf ANOVA p = 0.6682, 7 dpf ANOVA p = 0.4251). Sample sizes for all comparisons are the same as the corresponding quantifications in Fig. 1 for a-c and Fig. 2 for d.

**Supplementary Figure 4.**
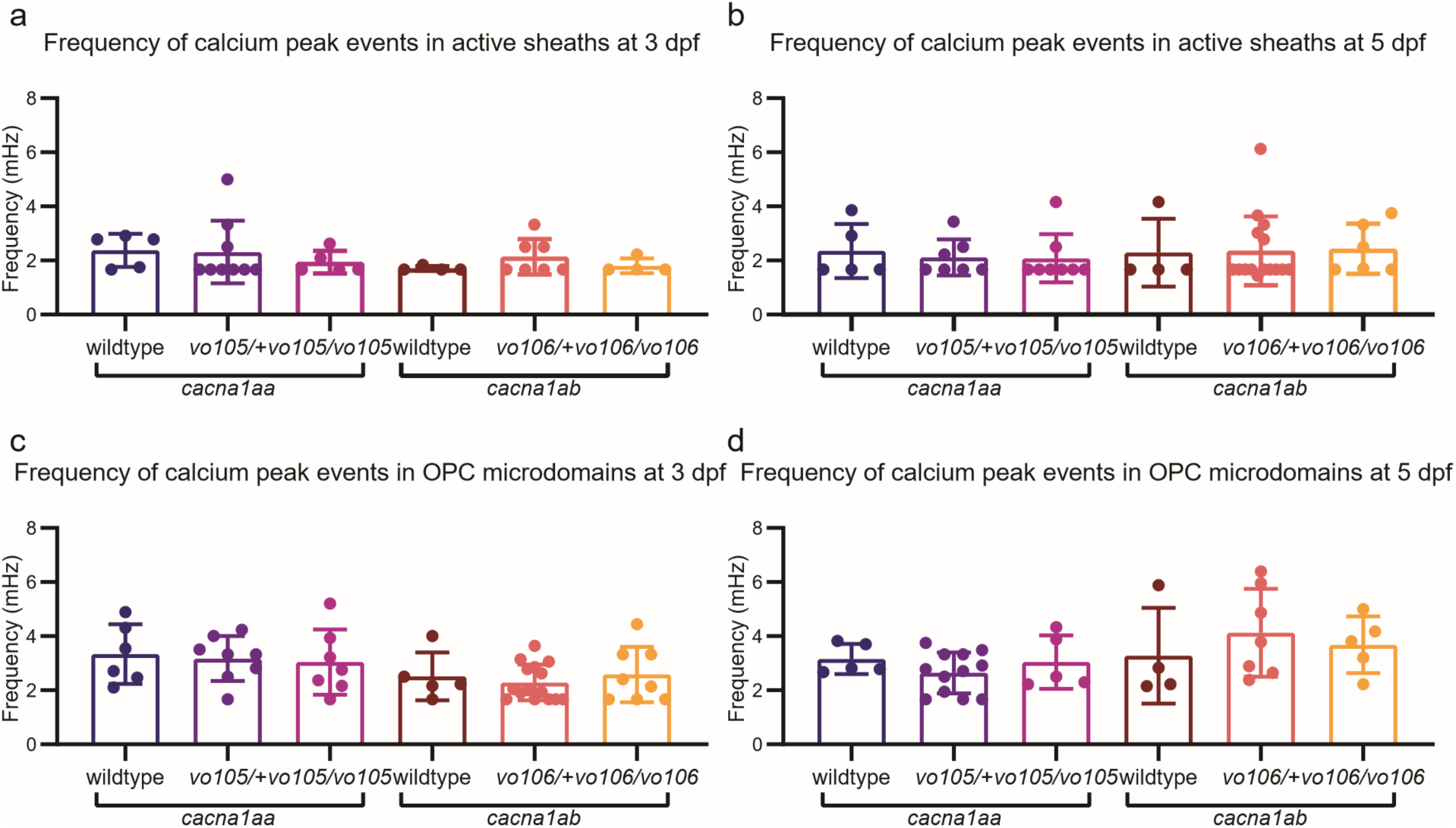
Calcium peak frequency in active myelin sheaths and OPC microdomains are similar across genotypes. Frequency of sheath calcium peak events, as seen in Fig. 3, is similar across genotypes both at a. 3 dpf (*cacna1aa* ANOVA p = 0.7024, *cacna1ab* ANOVA p = 0.3287), and at b. 5 dpf (*cacna1aa* ANOVA p = 0.8428, *cacna1ab* ANOVA p = 0.9819). Frequency of OPC microdomain calcium events, as seen in Fig. 4, is similar across genotypes both at c. 3 dpf (*cacna1aa* ANOVA p = 0.8820, *cacna1ab* ANOVA p = 0.6869) and at d. 5 dpf (*cacna1aa* ANOVA p = 0.3996, *cacna1ab* ANOVA p = 0.6640). Sample sizes for all comparisons are the same as corresponding quantifications in Fig. 3 for a-b and Fig. 4 for c-d.

**Supplementary Figure 5.**
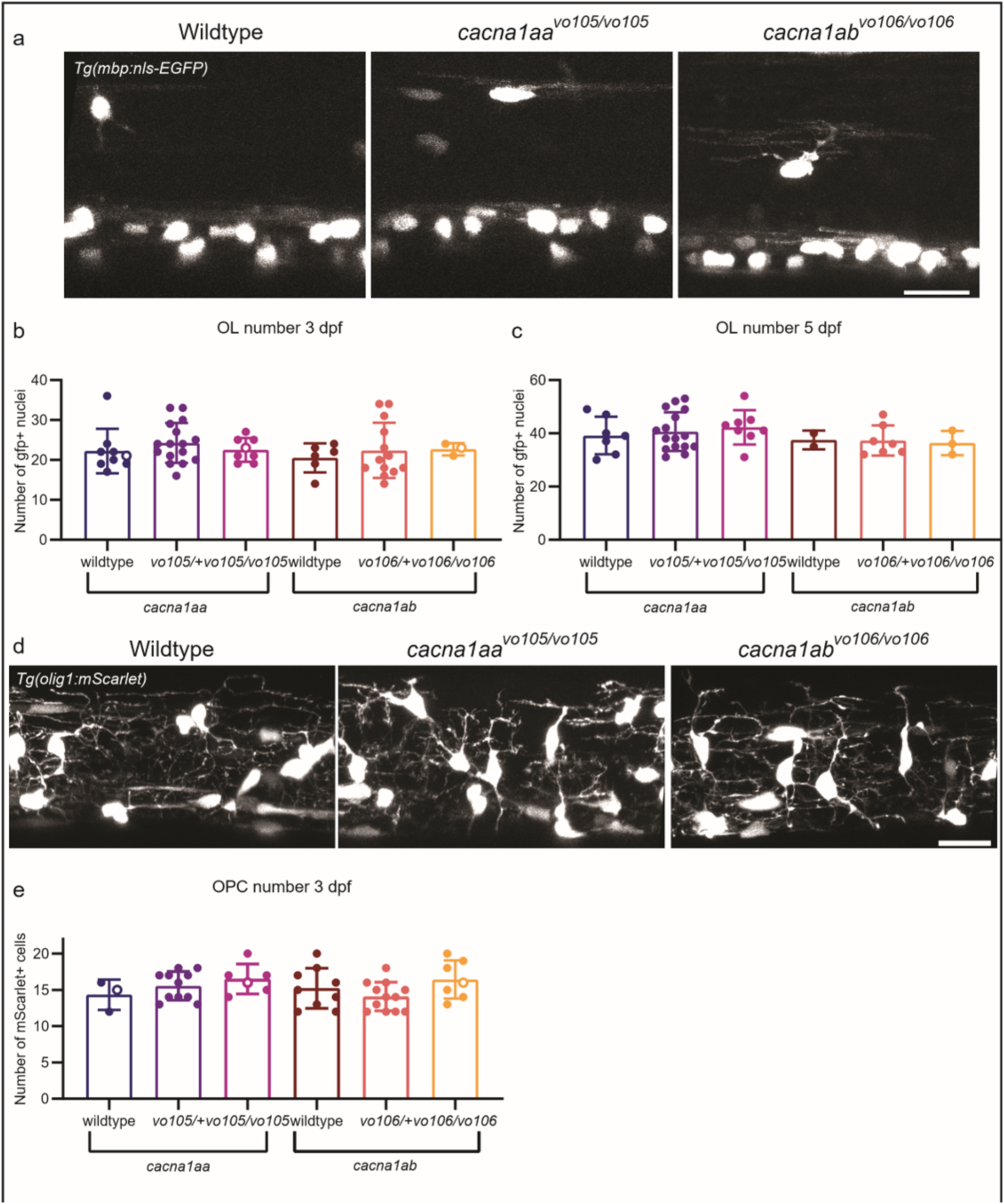
OLC numbers are similar in wild-type and P/Q-type channel mutant zebrafish. a. Example images from 3 dpf zebrafish spinal cord with *Tg(mbp:nls-EGFP)* in wild-type, *cacna1aa^vo105/105^*, and *cacna1ab^vo106/vo106^* zebrafish. b. Quantification of OLs at 3 dpf per body segment based on *mbp^+^* nuclei, as shown in a (*cacna1aa* ANOVA p = 0.5276, *cacna1ab* ANOVA p = 0.7847). Sample sizes are as follows: *cacna1aa* wildtype N = 9 fish, *cacna1aa^vo105/+^* N = 16 fish, *cacna1aa^vo105/vo105^* N = 8 fish, *cacna1ab* wildtype N = 6 fish, *cacna1ab^vo106/+^* N = 13 fish, *cacna1ab^vo106/vo106^* N = 3 fish. c. Quantification of OLs at 5 dpf per body segment based on *mbp^+^*nuclei (*cacna1aa* ANOVA p = 0.6958, *cacna1ab* ANOVA p = 0.9587). Sample sizes are as follows: *cacna1aa* wildtype N = 7 fish, *cacna1aa^vo105/+^* N = 16 fish, *cacna1aa^vo105/vo105^* N = 8 fish, *cacna1ab* wildtype N = 2 fish, *cacna1ab^vo106/+^* N = 7 fish, *cacna1ab^vo106/vo106^* N = 3 fish. d. Example images from 3 dpf zebrafish spinal cord with *Tg(olig1:mScarlet)* in wild-type, *cacna1aa^vo105/105^* and *cacna1ab^vo106/vo106^* zebrafish. e. Quantification of OPCs per body segment at 3 dpf, as shown in d (*cacna1aa* ANOVA p = 0.3279, *cacna1ab* ANOVA p = 0.1413). Sample sizes are as follows: *cacna1aa* wildtype N = 2 fish, *cacna1aa^vo105/+^*N = 11 fish, *cacna1aa^vo105/vo105^* N = 6 fish, *cacna1ab* wildtype N = 9 fish, *cacna1ab^vo106/+^*N = 12 fish, *cacna1ab^vo106/vo106^* N = 7 fish.

